# Gut microbiota of homing pigeons shows summer-winter variation under constant diet indicating a substantial effect of temperature

**DOI:** 10.1101/2022.05.18.492417

**Authors:** Maurine W. Dietz, Kevin D. Matson, Maaike A. Versteegh, Marco van der Velde, Henk K. Parmentier, Joop. A.J. Arts, Joana F. Salles, B. Irene Tieleman

## Abstract

**Background:** Gut microbiotas play a pivotal role in host physiology and behaviour, and may affect host life-history traits such as seasonal variation in host phenotypic state. Generally, seasonal gut microbiota variation is attributed to seasonal diet variation. However, seasonal temperature and day length variation may also drive gut microbiota variation. We investigated summer-winter differences in gut microbiota in 14 homing pigeons living outdoors under a constant diet by collecting cloacal swabs in both seasons during two years. Because temperature effects may be mediated by host metabolism, we determined basal metabolic rate (BMR) and body mass. Immune competence is influenced by day length and has a close relationship with gut microbiota, and it may thus be a link between day length and gut microbiota. Therefore, we measured seven innate immune indices. We expected gut microbiota to show summer-winter differences and gut microbiota to correlate with metabolism and immune indices.

**Results:** BMR, body mass, and two immune indices varied seasonally, other host factors did not. Gut microbiota showed differences between seasons and sexes, and correlated with metabolism and immune indices. The most abundant genus (*Lachnoclostridium 12*, 12%) and associated higher taxa, were more abundant in winter, though not significantly at the phylum level, *Firmicutes. Bacteroidetes* were more abundant in summer. The *Firmicutes*:*Bacteroidetes* ratio tended to be higher in winter. The KEGG ortholog functions for fatty acid biosynthesis and linoleic acid metabolism (PICRUSt2) had increased abundances in winter.

**Conclusions:** The gut microbiota of homing pigeons varied seasonally, even under a constant diet. The correlations between immune indices and gut microbiota did not involve consistently specific immune indices and included only one of the two immune indices that showed seasonal differences, suggesting that immune competence may be an unlikely link between day length and gut microbiota. The correlations between gut microbiota and metabolism indices, the higher *Firmicutes*:*Bacteroidetes* ratio in winter, and the resemblance of the summer-winter differences in gut microbiota with the general temperature effects on gut microbiota in the literature, suggest that temperature partly drove the summer-winter differences in gut microbiota in homing pigeons.

## Introduction

Animals anticipate and respond to changes in environmental conditions. A significant driver of environmental changes is seasonal variation in climate, which can be expected from highly predictable cues like day length. Seasonal abiotic factors are often associated with changes in different facets of organismal biology, such as reproductive and physiological state, food abundance, and behaviour. Animals may rely on predictable cues like day length, their endogenous annual pacemakers, or both to respond to predictable seasonal environmental variation [1,2]. As a result, the annual life cycles of animals, characterized by changes in behavioural, physiological, and morphological phenotypes [2–6], are predictably timed and maintained in captivity [5,7].

The microbiota (i.e., bacteria, archaea, lower and higher eukaryotes, and viruses [8]) living in and on individuals can also influence the physiology and behaviour of animals [9,10]. The gut microbiota is one of the largest animal microbial communities, and its symbiotic relationships with the hosts are often complex and bidirectional [10]. For example, the diet choice of hosts can strongly affect the composition and function of the hosts’ gut microbiota. At the same time, the gut microbiota can influence diet selection via the production of essential amino acids, such as tryptophan [9–12]. Because the gut microbiota influences host physiology and behaviour, the gut microbiota contributes to the phenotypic flexibility of their hosts [10] and thus may assist hosts in responding to seasonal changes. Hence, the gut microbiota is expected vary seasonally, as it does in some wild animals [13–16]. This seasonal variation in gut microbiota seems to be primarily driven by seasonal variation in diet [17]. In North American red squirrels (*Tamiasciurus husonicus*), for example, seasonal rhythms in the relative abundances of *Oscillospira* and *Corpococcus* genera were associated with seasonal variation in food availability [13]. While in the greater sage-grouse (*Centrocercus urophasianus*), seasonal variation in food quality likely explains the seasonal variation in gut microbiota composition and richness, possibly in combination with the seasonal variation in food- and water-associated microbiota [18].

In addition to diet, two seasonally varying abiotic factors may contribute to the seasonal variation in gut microbiota: temperature and day length [19–22]. Gut microbiota composition and function can change rapidly with changing temperature. For example, the gut microbiota of captive mice (*Mus musculus*) and Eastern red-backed salamanders (*Plethodon cinereus*) changed in microbial diversity, community composition, and relative abundances of different taxa after the animals were housed at low temperatures for 7-11 days [23–25]. In many host species, the effects of temperature change on gut microbiota are reflected in a variation in the relative abundances of different taxa. In vertebrates, *Firmicutes* are generally more abundant at lower ambient temperatures, while *Bacteroidetes* are more abundant at higher temperatures [20,23]. How ambient temperature directly affects gut microbiota is unclear, but mechanisms related to host metabolism are likely to play a role. For instance, temperature influences host metabolism via changing thermoregulation costs (independently of its gut microbiota), and the changing metabolism may shape the gut microbiota [19]. Transfers of gut microbiota (via co-housing and caecal or faecal transplantations) from hosts that were cold-acclimatized to those that were not induced changes in the metabolism and nutrient assimilation of the recipients. These physiological changes including the promotion of browning of white fat depots and the elevation of metabolic rate [20,23,24,26,27], mimicked changes due to cold-acclimatization. This suggests that seasonal temperature differences may result in a seasonal variation in gut microbiota that helps drive the seasonal acclimatization of hosts.

Seasonal variation in gut microbiota may also relate to variation in day length or the light-dark cycle. Day length variation affects the circadian clocks of animal hosts, leading to, for example, seasonal acclimatization [21,22,28]. The relationships between the host’s circadian systems and gut microbiotas are complex and bidirectional. The importance of the host circadian clock in maintaining circadian rhythms in gut microbiota is shown by the loss of the diurnal oscillations in the number of mucosal-resident bacteria in Per1/2 mice (*Mus musculus*) that lack their core molecular clock [29]. The gut microbiota’s circadian rhythm in such mice can be restored through time-restricted feeding [29,30]. Daily rhythms in hosts can also govern the effects of gut microbiota on other physiological systems of the hosts. For example, the host’s circadian system mediates the vital communication between gut microbiota and the host’s immune system [22]. Likewise, daily rhythms in gut microbiota can affect hosts: e.g., microbiota-produced short-chain fatty acids and bile acids can induce circadian entrainment in certain tissues and modulate hepatic circadian gene expression in mice [28]. Most research to date has focused narrowly on the direct effects of day length on the daily rhythm in gut microbiota characteristics. It remains unclear whether seasonal patterns in day length result in corresponding seasonal patterns in gut microbiota and whether this interaction aids hosts in adjusting to seasonal environmental variation.

Via host-mediated effects, seasonal variation in temperature and day length may contribute to the seasonal variation in gut microbiota, above and beyond any effect of seasonal changes in diet (Fig. 1). To explore this possibility, we investigated the summer-winter differences in gut microbiota in relation to host physiology in homing pigeons (*Columba livia*) that were fed a constant diet. During summer and winter of two consecutive years, we collected cloacal swabs and other host-related data from 14 individuals (six females and eight males) housed in outdoor aviaries. Because diet was constant, summer-winter differences in gut microbiota can be attributed to temperature and day length variation. As host metabolism may mediate the effects of temperature on gut microbiota, we investigated whether summer-winter variation in host metabolism was correlated with summer-winter variation in gut microbiota. To do so, we quantified basal metabolic rate (BMR) and body mass during both years, and daily food intake and digestive efficiency during summer and winter of the first year. In addition, we compared our results with the general effects of temperature variation on gut microbiota found in the literature. The link between day length and gut microbiota may be immune competence, because day length may affect the immune system and via the circadian clock also the communication between the immune system and the gut microbiota. Therefore, we assessed seven indices of the innate immune system during both years. We expected 1) that gut microbiota would show summer-winter differences despite a constant *ad libitum* food source; 2) that *Firmicutes* would be relatively more abundant in the winter due to lower temperatures, and *Bacteroidetes* would be relatively more abundant in summer cf. [20, 23]; 3) that metabolic differences would parallel summer-winter differences in gut microbiota since temperature affects host metabolism, and host metabolism influences the gut microbiota [19]; and 4) that immunological differences would parallel summer-winter differences in gut microbiota since day length may directly affect the immune system and mediate, via the host’s circadian clock, the communication between gut microbiota and the immune system [22].

**Fig. 1.**
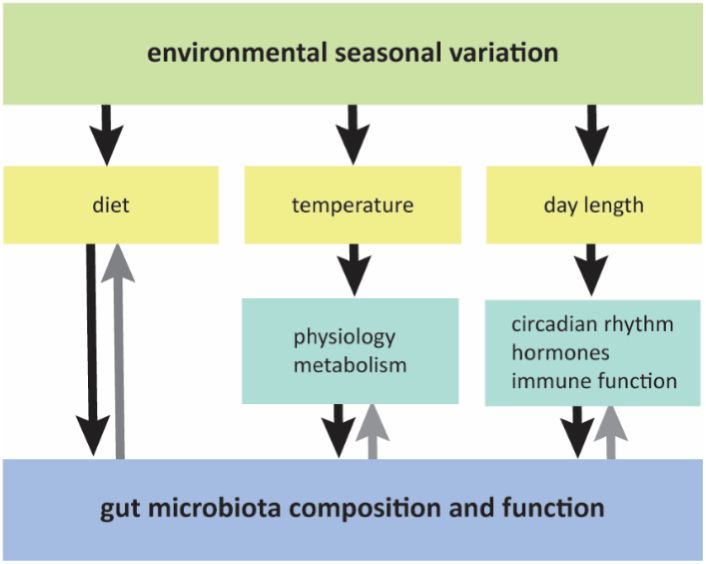
Schematic overview of direct and indirect pathways via which season may impact the gut microbiota. Yellow boxes indicate the three important aspects of seasonal environmental variation (diet, temperature, day length) that may impact the gut microbiota. Green boxes indicate the host components of the indirect pathways. E.g., seasonal day length variation impacts the circadian system of the host, which will impact gut microbiota composition and function via seasonal variation in feeding patterns and circadian rhythms in hormone expression. On its turn, the gut microbiota (blue box) impacts the host circadian system short chain fatty acids and bile acids production. Black arrows indicate the route of seasonal pathways modulating the gut microbiota, grey arrows indicate modulation of the host by the gut microbiota.

## Methods

### Animals

We used 14 homing pigeons (six females and eight males) hatched in captivity in late 2005, housed in same sex groups of 2–4 individuals in outdoor aviaries (4.01 m × 1.67 m × 2.2 m, l×w×h) at the Groningen Institute for Evolutionary Life Sciences (GELIFES) of the University of Groningen (N53°14.579’ E6°32.271’). Food (seed mixture 4 seasons for homing pigeons KASPER™ 6705, and pigeon pellets KASPER™ P40, Kasper Faunafood, Woerden, Netherlands; see supplementary information Table S1 for composition), grit and water were available *ad libitum*. All birds were exposed to outside air temperature and natural day length (see Table 1 for summer-winter differences in the experimental years). The birds were colour banded for individual identification.

**Table 1.**
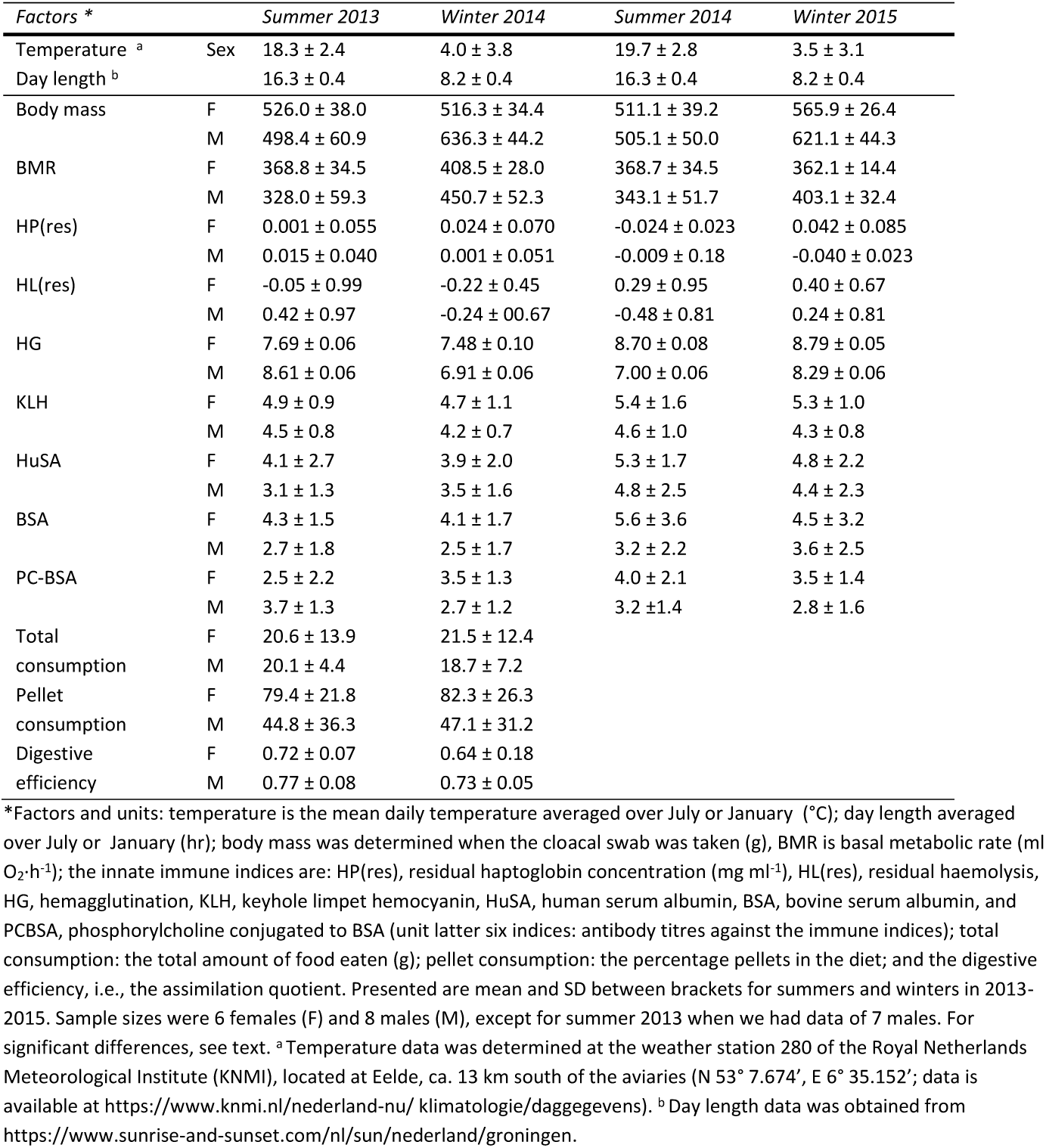
Seasonal variation in day length, temperature, body mass, BMR, immune indices, diet and digestive efficiency.

| Factors * |  | Summer 2013 | Winter 2014 | Summer 2014 | Winter 2015 |
| --- | --- | --- | --- | --- | --- |
| Temperature <sup>a</sup> | Sex | 18.3 ± 2.4 | 4.0 ± 3.8 | 19.7 ± 2.8 | 3.5 ± 3.1 |
| Day length <sup>b</sup> |  | 16.3 ± 0.4 | 8.2 ± 0.4 | 16.3 ± 0.4 | 8.2 ± 0.4 |
| Body mass | F | 526.0 ± 38.0 | 516.3 ± 34.4 | 511.1 ± 39.2 | 565.9 ± 26.4 |
|  | M | 498.4 ± 60.9 | 636.3 ± 44.2 | 505.1 ± 50.0 | 621.1 ± 44.3 |
| BMR | F | 368.8 ± 34.5 | 408.5 ± 28.0 | 368.7 ± 34.5 | 362.1 ± 14.4 |
|  | M | 328.0 ± 59.3 | 450.7 ± 52.3 | 343.1 ± 51.7 | 403.1 ± 32.4 |
| HP(res) | F | 0.001 ± 0.055 | 0.024 ± 0.070 | -0.024 ± 0.023 | 0.042 ± 0.085 |
|  | M | 0.015 ± 0.040 | 0.001 ± 0.051 | -0.009 ± 0.18 | -0.040 ± 0.023 |
| HL(res) | F | -0.05 ± 0.99 | -0.22 ± 0.45 | 0.29 ± 0.95 | 0.40 ± 0.67 |
|  | M | 0.42 ± 0.97 | -0.24 ± 0.67 | -0.48 ± 0.81 | 0.24 ± 0.81 |
| HG | F | 7.69 ± 0.06 | 7.48 ± 0.10 | 8.70 ± 0.08 | 8.79 ± 0.05 |
|  | M | 8.61 ± 0.06 | 6.91 ± 0.06 | 7.00 ± 0.06 | 8.29 ± 0.06 |
| KLH | F | 4.9 ± 0.9 | 4.7 ± 1.1 | 5.4 ± 1.6 | 5.3 ± 1.0 |
|  | M | 4.5 ± 0.8 | 4.2 ± 0.7 | 4.6 ± 1.0 | 4.3 ± 0.8 |
| HuSA | F | 4.1 ± 2.7 | 3.9 ± 2.0 | 5.3 ± 1.7 | 4.8 ± 2.2 |
|  | M | 3.1 ± 1.3 | 3.5 ± 1.6 | 4.8 ± 2.5 | 4.4 ± 2.3 |
| BSA | F | 4.3 ± 1.5 | 4.1 ± 1.7 | 5.6 ± 3.6 | 4.5 ± 3.2 |
|  | M | 2.7 ± 1.8 | 2.5 ± 1.7 | 3.2 ± 2.2 | 3.6 ± 2.5 |
| PC-BSA | F | 2.5 ± 2.2 | 3.5 ± 1.3 | 4.0 ± 2.1 | 3.5 ± 1.4 |
|  | M | 3.7 ± 1.3 | 2.7 ± 1.2 | 3.2 ± 1.4 | 2.8 ± 1.6 |
| Total consumption | F | 20.6 ± 13.9 | 21.5 ± 12.4 |  |  |
|  | M | 20.1 ± 4.4 | 18.7 ± 7.2 |  |  |
| Pellet consumption | F | 79.4 ± 21.8 | 82.3 ± 26.3 |  |  |
|  | M | 44.8 ± 36.3 | 47.1 ± 31.2 |  |  |
| Digestive efficiency | F | 0.72 ± 0.07 | 0.64 ± 0.18 |  |  |
|  | M | 0.77 ± 0.08 | 0.73 ± 0.05 |  |  |

\*Factors and units: temperature is the mean daily temperature averaged over July or January (°C); day length averaged over July or January (hr); body mass was determined when the cloacal swab was taken (g), BMR is basal metabolic rate ( $\text{ml O}_2 \cdot \text{h}^{-1}$ ); the innate immune indices are: HP(res), residual haptoglobin concentration ( $\text{mg ml}^{-1}$ ), HL(res), residual haemolysis, HG, hemagglutination, KLH, keyhole limpet hemocyanin, HuSA, human serum albumin, BSA, bovine serum albumin, and PCBSA, phosphorylcholine conjugated to BSA (unit latter six indices: antibody titres against the immune indices); total consumption: the total amount of food eaten (g); pellet consumption: the percentage pellets in the diet; and the digestive efficiency, i.e., the assimilation quotient. Presented are mean and SD between brackets for summers and winters in 2013-2015. Sample sizes were 6 females (F) and 8 males (M), except for summer 2013 when we had data of 7 males. For significant differences, see text. <sup>a</sup> Temperature data was determined at the weather station 280 of the Royal Netherlands Meteorological Institute (KNMI), located at Eelde, ca. 13 km south of the aviaries (N 53° 7.674', E 6° 35.152'; data is available at https://www.knmi.nl/nederland-nu/klimatologie/daggegevens). <sup>b</sup> Day length data was obtained from https://www.sunrise-and-sunset.com/nl/sun/nederland/groningen.

### Cloacal swab collection

We collected cloacal swabs in July (summer) and January (winter) between 9:00-11:00 CET for two consecutive years, starting in July 2013. We inserted a sterile viscose swab into the cloaca without contacting feathers or skin and gently rotated it for 10 s in the intestinal lumen. Swab tips, which were cut from the shaft using scissors sterilized with 76% ethanol then flamed, were stored in a sterilized 1.5 ml vials. After adding a drop of sterile PBS, swabs were stored at -20°C until analysis. We randomized the sampling order of the individual pigeons per season and sampled two pigeons per day (rate limited by further handling protocols describes below). Body mass was recorded after swab collection (± 0.1 g).

### Daily food intake and digestive efficiency

We determined daily food intake and digestive efficiency after cloacal sampling, and placed for this the pigeons individually in clean outdoor aviaries located in the same aviary cluster as their home aviary. Each bird was offered ∼50 g of pellets, ∼60 g of wheat, ∼30 g of corn, and ∼30 g of green peas (the latter three are the main seeds provided and eaten from the seed mixture offered), and *ad libitum* water. The next day between 11:00-13:00 CET, we removed the birds from the aviaries, recorded body mass again, and transferred them indoors for the basal metabolic rate measurement. All faeces and food leftovers were collected and weighed. Faeces were stored at -20°C until analysis.

We determined the energy and water contents of the food items (pellets, wheat, corn, and green peas) once for each food item and used these data to calculate dry food mass eaten and energy intake for each food item per trial. Faecal energy content was determined per individual trial. Before determining energy contents, we dried the food items and faeces to constant mass at 60°C, i.e., until the change between weightings was <0.1 % of the initial fresh mass (all masses ±0.0001 g). This took ∼13 d for food items and ∼6 d for faeces. We ground dry food and faeces to powder (Retsch grinder ZM 100), pressed them into pills (∼1 g), and dried them to constant mass at 60°C to determine pill dry mass (±0.0001 g). We burned the pills in an adiabatic bomb calorimeter (IKA C 5000) to determine their energy content (kJ·g^-1^). We analysed all samples at least in duplicate, which in general differed by <2 % of the lower energy content pill. Two samples were measured in triplicate. The mean energy content of the replicates was used in further analyses. For each trial, the digestive efficiency or assimilation quotient was calculated as:

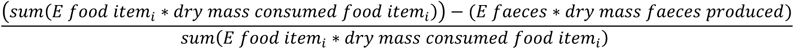

where *E* is the energy content of the dry *food item*_*i*_ or faeces, and *food item*_*i*_ is pellets, wheat, corn, or green peas.

During the first year, pigeons ate typically > 20 g food (summer 2013 23.16 ± 10.28 g, winter 2014 20.09 ± 12.15 g), while in the second year, almost all pigeons ate much less (summer 2014 7.88 ± 6.23 g, winter 2015 5.51 ± 7.93 g). Hence, we disregarded the second year’s data. The lower food consumption in the second year was unexpected, and we cannot explain this observation as generally repeating procedures with animals is expected to result in less stress. In the second year, we did determine the food intake of a few pigeons left isolated in their home cage. Food amounts eaten by these birds were similar to the 2013-2014 values, suggesting that not isolation itself but rather the aviary transfer in combination with isolation led to the lower food consumption in the second year.

### Basal metabolic rate

Prior to measuring BMR, the pigeons were placed individually in a darkened box (30 cm × 25 cm × 28 cm) indoors to acclimatize and fast for ∼4 h. At ∼17:00 CET, the birds were placed individually into 13.5 l metabolic chambers and placed inside a climatic chamber set at 25 ± 0.5°C (thermoneutral for domestic pigeons [31,32]). Oxygen consumption was measured throughout the night using standard flow-through respirometry methods and recorded during 17-minute windows alternately for each individual (for details, see [33]). The following day at ∼ 9:00 CET, the bird was removed from the metabolic chamber and returned to its aviary. Body mass was recorded immediately before and after the measurement. BMR (ml O_2_·h^-1^) was based on the lowest average oxygen consumption during any of the 17-minute windows recorded throughout the night.

### Immune indices

We assessed innate immune competence from blood samples collected ca. one month prior to each microbiota sampling moment. We measured seven innate immune indices using three assay types. First, we used a commercially available colorimetric assay (TP801; Tri-Delta Diagnostics, NJ, USA) to quantify haptoglobin concentration (or its haem-binding functional equivalents, mg·ml^-1^). We followed the manufacturer’s instructions with the additions and changes described in [34]. Because this functional assay is sensitive to contamination by haem in haemolysed samples, we measured sample redness (absorbance at 450nm), a proxy for haemolysis, prior to the addition of the second reagent and the initiation of the colour change reaction [34]. In the current dataset, the relationship between sample redness and haptoglobin was significant (*P* = 0.02), so we used the residual variation in haptoglobin in further analyses. Haptoglobin did not vary with sample age.

Second, we used a haemolysis-haemagglutination assay to measure titres of complement-mediated lysis, and natural antibody- (NAb-) mediated agglutination of rabbit erythrocytes [35]. Agglutination was recorded from plate images made 20 min post-incubation; lysis was recorded from plate images made 24 h after incubation, as described in [36]. In the current dataset, the relationship between sample age and lysis (but not sample age and agglutination) was significant (agglutination, *P* = 0.82; lysis, *P* < 0.01); hence we used the residual variation in lysis in further analyses.

Third, we used indirect three step ELISA to measure titres of NAbs against four antigens separately, none of which individuals had been previously vaccinated against: keyhole limpet hemocyanin (KLH), human serum albumin (HuSA), bovine serum albumin (BSA), and phosphorylcholine conjugated to BSA (PC-BSA) [37]. In brief, wells were incubated with 100uL of coating buffer (pH 9.6) containing one of the four antigens for one hour at 37°C. Wells were then washed (water + 0.05% Tween 20), blocked (phosphate buffer saline (PBS) + 1% horse serum + 0.05% Tween 20) for 30 minutes, and washed again. Plasma samples were serially four step diluted (KLH: 1:40, 1:160, 1:640, 1:2560; other antigens two step dilutions: 1:20, 1:40, 1:80, 1:160) in wells containing 100uL dilution buffer (PBS + 0.5% horse serum + 0.05% Tween 20). Duplicate standard positive plasma samples (a pool of pigeons) were two step diluted with dilution buffer. Two antibodies were added and incubated (1hr at 37°C) sequentially: first, 100μL of a 1:5000 dilution of rabbit-anti-pigeon antibodies (IgG(H+L); Nordic; batch no. 6162); second, 100μL of a 1:2000 swine-anti-rabbit antibodies conjugated to horseradish peroxidase. After each incubation, wells were washed. The colour change reaction was initiated with the addition of substrate (containing reverse osmosis purified water, 10% tetramethylbenzidine buffer [15.0 g/L sodium acetate, and 1.43 g/L urea hydrogen peroxide; pH 5.5], and 1% tetramethylbenziding [8 g/L TMB in DMSO]) at room temperature and stopped (with 50μl/well of 1.25M H_2_SO_4_) after 15 minutes; absorbance was read at 450nm with a Multiskan Go (Thermo scientific). All titres of the NAbs were not correlated with sample age (all *P* > 0.1). Antibody titres were calculated as described by [38] (taken from [39]). For details on the antibody titres calculation see the Supporting Information.

### DNA isolation and 16S rRNA gene amplicon sequencing

We randomized the cloacal swabs prior to DNA extraction. DNA was isolated from the samples using the FastDNA™ kit for Soil (MP Biomedicals, Santa Ana, CA, USA) according to the manufacturer’s instructions. Two exceptions were cell lysis, which was achieved by beat-beating three times one minute instead of three minutes continuously, to prevent the samples from heating up, and the DNA elution which was done using in 100 μl PCR-grade water. We quantified sample DNA concentrations using the Quant-it PicoGreen dsDNA kit (Molecular Probes, Invitrogen, Eugene, OR, USA) and normalized the DNA concentrations in the subsequent PCR to 1 ng template DNA per 25 μl reaction. The samples were randomized again before amplifying the V4/V5 region of the 16S rRNA gene in a triplicate using the primers 515F and 926R [40,41] with Illumina adaptors at the 5’-end. We used the following thermal cycling protocol: 5 min at 95°C, 35 cycles with 40 s at 95°C, 45 s at 56°C, 40 s at 72°C, followed by 10 min at 72°C. We pooled the triplicates after the PCR. We excluded one cloacal sample with poor PCR results (a male, summer 2013), and sent after purification (QIAquick gel extraction Kit, QIAGEN GmbH, Hilden, Germany) the 55 pigeon samples, a negative control swab and 4 negative PCR controls, to GenoToul (INRA, Toulouse, France) for library preparations and Illumina sequencing using 2 × 250 bp v2 chemistry. At GenoToul, the sequence reads were demultiplexed and quality filtered using the default settings in QIIME.

### Sequence data processing

We processed the raw sequence data using the standard QIIME2 protocol (v2018.2 [42]). Using the DADA2 (v2018.2) pipeline, we trimmed the primers, truncated and merged the forward and reverse reads based on quality plots, and removed chimera. The taxonomy table was built using the Silva v132 reference database [43,44]. Next, we filtered *Archaea*, chloroplasts, mitochondria, and vertebrates from the data. The end products, an Amplicon Sequence Variant (ASV) table and the phylogenetic tree were further processed in R (v4.0.2 [45]) using *Phyloseq* (v1.32.0 [46]) and *vegan* (v2.5-6 [47]). At this stage, the data included 1,056 taxa, and the total number of sequence reads was 1,606,609, with counts ranging between 1,779-95,837 reads for cloacal swab samples and between 78-1,449 reads for the negative controls.

### Statistical analysis

#### Host parameters

We used linear mixed models (LMM, *nlme* v3.1-148 [50]) to identify correlations between the host parameters (metabolic and immune indices) and season (summer vs. winter), sex, their interaction term (fixed factors), and individual bird colour bands (BirdID) nested within aviary (random factors), using a stepwise backward exclusion of nonsignificant fixed factors. At each step of the analysis, the normality of the model and homoskedasticity of the residuals were checked. For the final model, we tested if aviary as a random factor contributed significantly (ANOVA), which was never the case. Hence the final model contained only BirdID as random factor.

#### Contamination

We used *Decontam* (v1.8.0 [48]) to identify general contaminants using the recommended settings. The *Decontam* frequency method identified eight out of the 1,056 ASVs as contaminants. However, two of the eight were not present in the negative controls indicating that the identification of contaminants using this method was unreliable. The *Decontam* prevalence method identified six general contaminants, but three of the six was had a prevalence of only one in the cloacal swab samples, again a sign of unreliability of the method. Given these results, and considering the very low read counts of the negative controls (78–1,449 reads), we concluded that no contaminants with a considerable impact on the data could be detected. We therefore removed the reads from negative control samples from the data set. Hereafter the data included 1,032 taxa divided over 55 samples, with in total 1,602,954 reads.

We checked the data for rare ASVs based on read counts and prevalence. Since we used DADA2 to trim primers and merge and truncate primers, the data initially contained no singletons. After removing the negative control samples, there were 5 singleton ASVs (0.5%) and 185 doubleton ASVs (17.9%), indicating that only few ASVs had low read counts. Prevalence analysis showed that despite having four samples per individual, 81.3% of the ASVs occurred in only 1 sample, indicating a high variability between the samples within and among individuals. This percentage is comparable to other data sets of ours, including a dataset from captive juvenile rock pigeons that includes eight samples per individual (77.8%) and a dataset from adult free-living feral pigeons (61.3%) (Dietz pers. comm.). These percentages are also comparable to those from a variety of other species [49].

Because richness rarefactions curves levelled off around 3,000 reads (see Supplementary Information Fig. S1), we rarefied the data to 3,270 reads, which equalled to reads of the sample with the second lowest number of reads. Rarefying eliminated thus one sample (a male, winter 2015, 1,779 reads) from the data set. Thereafter, 598 taxa were left divided over 54 samples (for females six samples per season, and for males seven samples in summer 2013 and winter 2015, and eight samples in winter 2014 and summer 2014). The percentage of singletons increased after rarefying to 37.6%, while 13.5% of the ASVs were doubletons. The percentage of very low prevalence ASVs remained comparable to before rarefying, with 71.2% of the ASVs occurring in one sample.

#### Alpha-diversity

Similar to the host parameters, we identified correlations between alpha-diversity indices (richness, Shannon index, and Faith’s phylogenetic diversity) and season (summer vs. winter), sex, their interaction term (fixed factors) using linear mixed models with BirdID nested within aviary as random factors. We used again a stepwise backward exclusion of nonsignificant fixed factors and checked at each step of the analysis the normality of the model and homoskedasticity of the residuals. For the final model, we tested if aviary as a random factor contributed significantly (ANOVA). Using the same procedure, we next tested if the alpha-diversity indices were correlated with temperature or day length-related host characteristics in separate LMMs. To test temperature-related mechanisms, we used LMMs with metabolism indices (BMR and body mass) as fixed factors, and individual bird colour bands (BirdID) nested within aviary (random factors). Food intake and digestive efficiency were not included because they did not show seasonal variation (see Results) and because the data was limited to one year. To test day length-related mechanisms, we used LMMs with the seven innate immune indices as fixed factors, and individual bird colour bands (BirdID) nested within aviary (random factors). In almost all final models, aviary did not contribute significantly and was thus not included in the final models unless stated otherwise.

#### Bacterial community composition

The bacterial community composition (beta-diversity) was assessed by looking at the taxonomic similarities between seasons (summer vs winter) and sexes using the Jaccard similarity index (community membership: presence/absence), Bray-Curtis dissimilarities (community structure: presence/absence and abundance matrix), and by looking at the phylogenetic similarities between seasons and sexes using unweighted (community membership: presence/absence table) and weighted UniFrac distances (community structure: presence/absence/abundance matrix [51]). A principal coordinate ordination analysis (PCoA) of the beta-diversity indices was performed to test if community clustering and group dispersion differed between seasons or sexes, which was achieved by modelling beta-diversity (dis)similarities and distances from an ASV-level table using PERMANOVA with 10,000 permutations (adonis2 function in *vegan*) [52,53]. Since we had multiple samples per individual, we first evaluated the effect of individuals on the different beta-diversity indices; this effect was always significant (*P* < 0.01). We next tested for the effect of season, sex, and their interaction while including individual as a blocking factor (*strata*) to control for the repeated sampling. We evaluated the degree of within-group dispersions (permutest) using the ‘betadisper’ function [54] in *vegan*. These were always nonsignificant, indicating that differences found were not due to differences in group dispersions.

For the three beta-diversity indices that showed seasonal differences, we tested if their ordination was comparable to the ordination of the metabolism indices (BMR and body mass) or the immune indices by performing a Procrustes analysis using the *Procrustes* and *Protest* functions in *vegan* [52,53]. We analysed the similarity of the two-dimensional shapes produced from overlaying the principal component analyses of the Euclidian distances of metabolism or immune competence with the beta-diversity measure.

#### Taxonomic composition

We used the same LMM procedure as described above for the alpha-diversity indices to examine variation in the relative abundances of the most abundant phyla (>5%) and genera (>5%). As explanatory variables, we included season, sex, and their interaction term, metabolism (BMR and body mass), or immune competence (seven innate immune indices). Before running the LMMs, taxa proportions were logit transformed as log[(*p*+*e*)/(1-*p*+*e*)], where *p* is the proportion of a taxon in a given sample and *e* the lowest proportion (among samples) for that taxon excluding zero [55].

#### Bacterial biomarkers

To pinpoint which taxa may play a role in seasonal acclimatization in homing pigeons, we identified seasonal bacterial biomarkers via two methods. First, we characterized bacterial biomarkers as the ASVs that were more abundant in a season via a linear discriminant analysis (LDA) effect size analysis (LEfSe, [56]) on the online Huttenhower platform https://huttenhower.sph.harvard.edu/galaxy/), using the default settings. Since the Silva v132 database characterizes ASVs only at the genus level, we assigned a unique number to each ASV at the species level before performing the LEfSe analysis. The analysis was done for each sex separately, as sex significantly affected the indices of alpha- and beta-diversity.

Second, we characterized bacterial biomarkers per season and per sex based on prevalence by comparing the cores of a season or a sex with the overall core for all samples. ASVs were considered belonging to the core when present in 90% of the samples a group (*microbiome* v1.10.0 [57]). Hence, when all samples are taken into account, core ASVs should occur in at least 49 of the 54 samples. Core-based bacterial biomarkers for a season or sex where those ASVs that were unique for a season or sex when comparing their cores with the core of all samples.

#### Functional profile of gut microbiota

Lastly, we explored if the functional profile of the gut microbiota showed seasonal differences. We used PICRUSt2 (version 2.3.0b) to predict KEGG ortholog (KO) metagenome functions from the 16S rRNA gene data using the rarefied data set [58,59]. We tested with PERMANOVA if KO function abundances varied between summer and winter, and between males and females, following the same approach as for the beta-diversity indices. Individual was again significant (*P* < 0.01) and included as blocking factor to control for repeated sampling (*strata*). Next, we identified which KO functions were more abundant in summer and which were more abundant in winter using a LEfSe analysis [56] for each sex separately.

## Results

### Seasonal variation in host parameters

As is common among birds living in temperate areas, pigeon body mass was higher in winter than in summer (Table 1; LMM, season *P* = 0.05, sex *P* = 0.37, season*sex *P* < 0.01). Although the birds were heavier in winter, their total daily food intake did not differ between winter and summer, nor between sexes (LMM, season *P* = 0.89, sex *P* = 0.96, season*sex *P* = 0.75). The digestive efficiency also did not vary with season or sex (LMM, season *P* = 0.34, sex *P* = 0.28, season*sex *P* = 0.81).

Consumption of different food components (pellets or seeds) did not differ between seasons (percentage pellets eaten; LLM, season*sex *P* = 0.60, season *P* = 0.82). However, the average percentage pellets eaten by females (80.9%) was 1.8 times higher than the average percentage pellets eaten by males (46.0%; LMM, sex *P* = 0.02). The sexual variation in food preference had no implications for energy intake because the energy content did not differ between pellets and seeds (pellets: 17.47 kJ·g^-1^, corn: 18.11 kJ·g^-1^, peas: 17.89 kJ·g^-1^, wheat: 18.00 kJ·g^-1^). Hence all pigeons consumed the same amount of food and energy in summer and winter, and this did not differ between the sexes despite the sexual differences in diet preferences.

In general, BMR was higher in winter than in summer (Table 1; LMM, season *P* = 0.12, sex *P* = 0.18, season*sex *P* < 0.01). In both sexes, BMR was lower in winter 2015 than in winter 2014 (but in males, in both winters BMR was higher than either summer), despite similar winter temperatures (4.0°C in winter 2014 and 3.5°C in winter 2015, Table 1).

Titres of antibodies against phosphorylcholine conjugated to BSA (i.e., anti-PC-BSA) were higher in summer than in winter (Table 1; LMM, season *P* = 0.43, sex *P* = 0.53, season*sex *P* = 0.05). Haptoglobin concentrations (residuals after correcting for redness) varied with season and season*sex (Table 1; LMM, season *P* = 0.04, sex *P* = 0.53, season*sex *P* = 0.03). Haptoglobin concentrations corrected for redness were higher in females, and in females also higher in winter than in summer. The five other immune indices did not vary with sex, and unexpectedly, also did not vary with season or season*sex (LMM, all *P* > 0.05).

### Alpha-diversity

Richness was higher in females in winter than in summer but did not differ between seasons in males (season*sex *P* = 0.04, Table 2, Fig. 2a). In addition, richness was negatively correlated with body mass (Fig. 2d) but was not correlated with any immune index. Shannon diversity did not differ between summer and winter or sexes (Fig. 2b), nor was it correlated with any metabolism or immune index. Faith’s phylogenetic diversity was lower in males in winter but did not differ between seasons in females (season*sex *P* = 0.01, Table 2, Fig. 2c). Similar to the richness, Faith’s phylogenetic diversity was negatively correlated with body mass, but in addition, it was positively correlated with antibody titres against KLH (Fig. 2d, e).

**Table 2.** LMM analysis of alpha diversity indices ^a^.

| *Alpha diversity indices* | *Predictors final model* | *Df* | *F* | *P* |
| --- | --- | --- | --- | --- |
| Richness |  |  |  |  |
| vs. season, sex, season*sex | Season | 1,38 | 1.86 | 0.18 |
|  | Sex | 1,12 | 0.04 | 0.84 |
|  | Season*Sex | 1,38 | 4.35 | 0.04 |
| vs BMR and body mass | Body Mass | 1,39 | 7.92 | 0.01 |
| Faith's PD |  |  |  |  |
| vs season, sex, season*sex | Season | 1,38 | 1.36 | 0.25 |
|  | Sex | 1,12 | 0.08 | 0.78 |
|  | Season*Sex | 1,38 | 9.09 | 0.01 |
| vs BMR and body mass | Body mass | 1,39 | 14.22 | <0.01 |
| vs immune indices | KLH | 1,36 | 7.21 | 0.01 |

<sup>a</sup> Only the fixed factors of the final models are presented. Faith's PD is Faith's phylogenetic diversity.

**Fig. 2.**
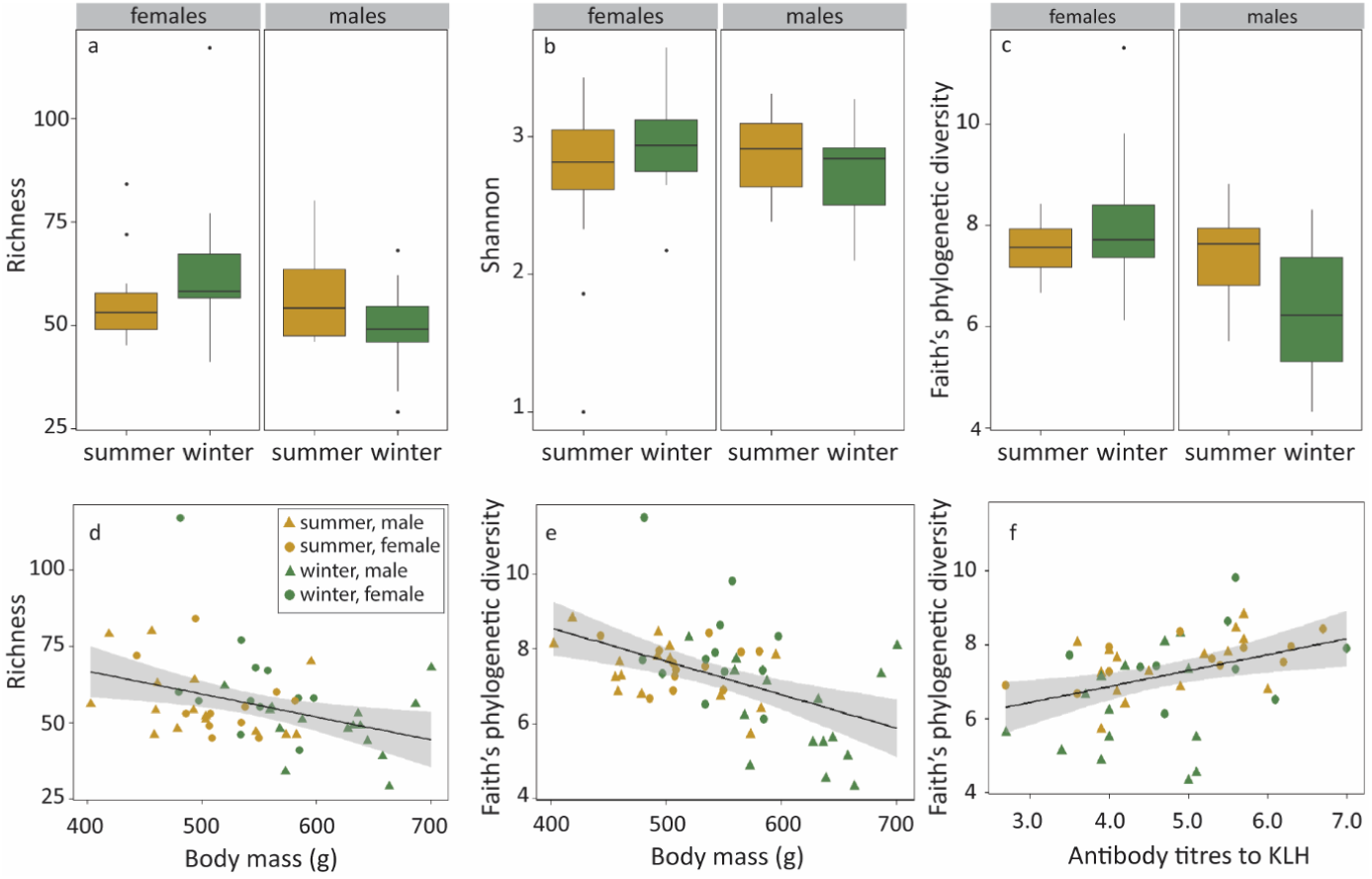
Relationships between alpha-diversity indices and season and sex, and metabolism and immune indices. The boxplots present seasonal and sexual variation for (**a**) richness, (**b**) Shannon diversity, and (**c**) Faith’s phylogenetic diversity. Richness and Faith’s phylogenetic diversity both decreased with increasing body mass (**d, e**), while Faith’s phylogenetic diversity also increased with titres of antibodies against KLH (**f**). Statistics are presented in Table 2.

### Bacterial community composition

Both taxon presence/absence (Jaccard, Bray-Curtis; Fig 3a, b) and phylogenetic (weighted UniFrac, Fig. 3d) community composition varied with season and sex, but not their interaction term (Table 3). Unweighted UniFrac did not vary with season or sex (Fig. 3c). Season explained less of the variation in community composition (2.6-3.1%) than sex (6.5-12.8%). The ordination of the Jaccard and Bray-Curtis (dis)similarities, and weighted UniFrac distances matched with that of the metabolism indices (BMR and body mass; all Procrustes SS = 0.88, *P* = 0.01, Fig. S2), but not with that of the immune indices (*P* = 0.96, *P* = 0.99, *P* = 0.73, for Jaccard, Bray-Curtis and weighted UniFrac, respectively).

**Table 3.** Permanova analyses of the beta diversity indices ^a^.

| Beta diversity indices | Predictors final model | R<sup>2</sup> (%) | F | P |
| --- | --- | --- | --- | --- |
| Jaccard | Season | 2.6 | 1.46 | <0.01 |
|  | Sex | 6.5 | 3.65 | <0.01 |
| Bray-Curtis | Season | 3.1 | 1.80 | <0.01 |
|  | Sex | 8.4 | 4.81 | <0.01 |
| Weighted UniFrac | Season | 2.6 | 1.55 | 0.01 |
|  | Sex | 12.8 | 7.71 | 0.01 |

<sup>a</sup> Only the final models are presented.

**Fig. 3.**
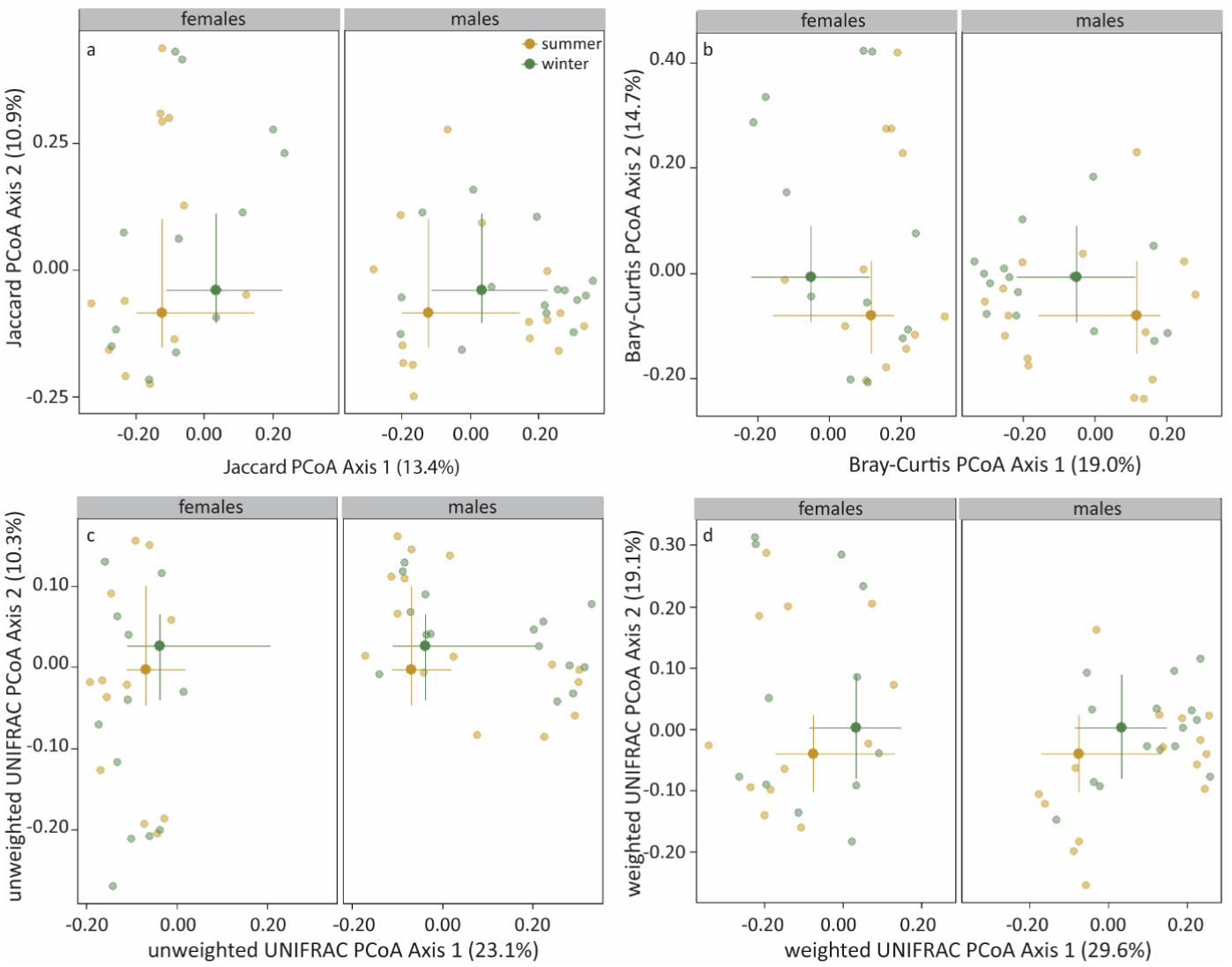
Seasonal and sexual variation in Jaccard (**a**) and Bray-Curtis (dis)similarities (**b**), and unweighted (**c**) and weighted UniFrac distances (**d**) depicted in PCoA plots. Statistics are presented in the text. The large symbols represent the medians, the error bars the 25% and 75% quantiles. The transparent symbols present the underlying data.

### Taxonomic composition

The taxa were divided over 14 phyla, including an unclassified phylum belonging to an unclassified kingdom. For the subsequent analyses, we divided the large *Proteobacteria* phylum into three classes: *Alphaproteobacteria, Betaproteobacteria*, and *Gammaproteobacteria*. Four phyla and a *Proteobacterium* class had high relative abundances (>5%), namely *Firmicutes* (43.1%), *Actinobacteria* (30.2%), *Fusobacteria* (10.3%), *Bacteroidetes* (8.2%), and *Gammaproteobacteria* (7.8%; Fig. 4). All of which, apart from the *Fusobacteria*, are commonly found in avian gut microbiota [60,61]. Of the most abundant phyla/*Proteobacteria* class, relative abundances of *Bacteroidetes* differed between seasons, with higher abundances in summer than in winter (*P* < 0.01, Fig. 4b, Table 4), while relative abundances of *Firmicutes* tended to differ between seasons (*P* = 0.07), with higher abundances in winter than in summer. Relative abundances of these two phyla and *Actinobacteria* and *Fusobacteria* also differed between sexes (Table 4). In addition, we found correlations with metabolism and immune indices. The logit proportion of *Bacteroidetes* was negatively correlated with BMR (*P* < 0.01, Fig 4c). The logit proportion of *Firmicutes* was positively correlated with body mass and negatively correlated antibody titres against BSA (*P* = 0.04 and *P* = 0.03, respectively, Fig 4d, e). The logit proportion of *Fusobacteria* was negatively correlated with haemolysis titres that had been corrected for sample age (*P* = 0.03, Fig. 4f). Lastly, logit proportions of *Gammaproteobacteria* did not show any differences between seasons or sexes but was positively correlated with antibody titres against BSA (*P* <0.01).

**Table 4.**
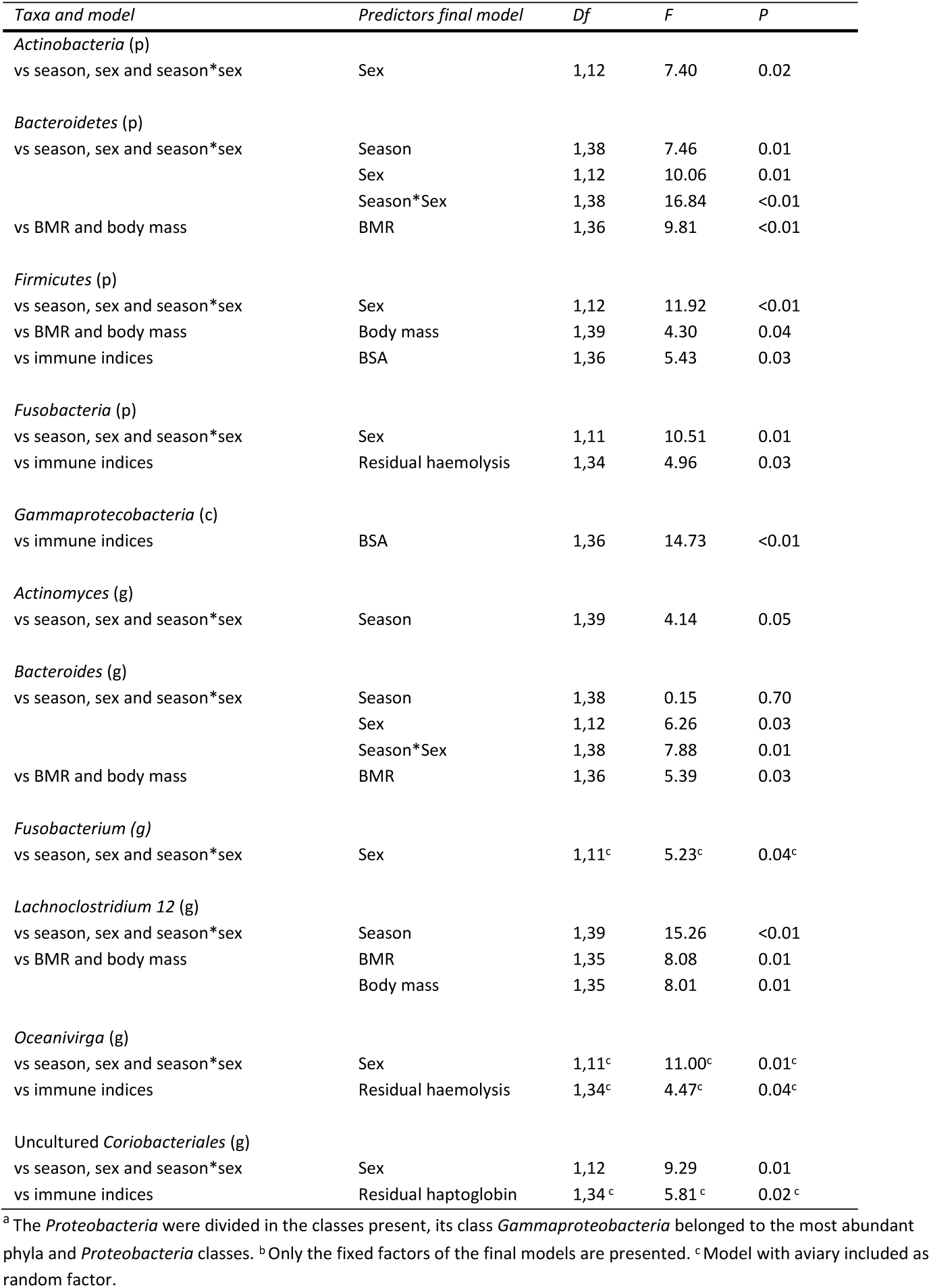
LMM analysis of the logit proportions of the most abundant phyla ^a^ and genera ^b^.

**Fig. 4.**
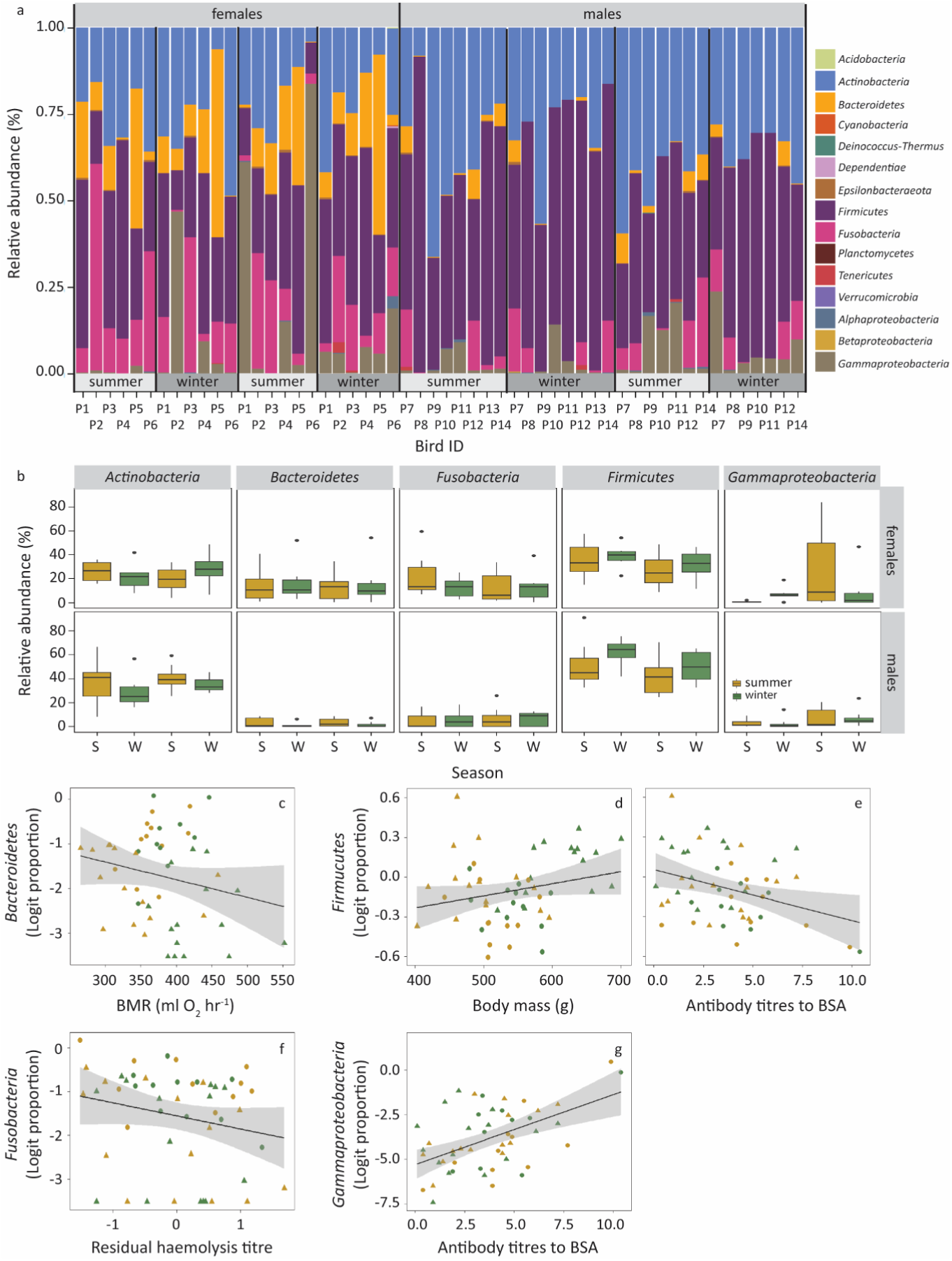
The variation in phylum relative abundance with season, sex, and immune indices. (**a**) Stacked bar plots of phylum relative abundance per sample. Note that the large phylum *Proteobacteria* was divided into the three classes present. (**b**) Boxplots of the relative abundances of the five most abundant phyla. The data is organized per sex and season (S = summer, W = winter), starting with the summer of 2013. The relationships between the logit(proportion) of *Bacteroidetes* and BMR (**c**), the logit(proportion) of *Firmicutes* and body mass (**d**) and BSA titre (**e**), the logit(proportion) of *Fusobacteria* and residuals of the haemolysis titre (**f**), and the logit(proportion) of *Gammaproteobacteria* and BSA titre (**g**) are presented in separate panels. Statistics are presented in Table 4.

Eight of the 128 genera present had high relative abundances (>5%). *Lachnoclostridium 12* (*Firmicutes*) was the most abundant genus (12.9%) due to one ASV that was the most abundant ASV and present in all samples (ASV nr. 093e1dd8072a68e5fa46226677183da). Similar to its phylum, the logit proportion of *Lachnoclostridium 12* was higher in winter than in summer, but now significantly (*P* < 0.01). The logit proportion of *Lachnoclostridium 12* did not vary with sex, was slightly negatively correlated with BMR, and positively correlated with body mass (Fig. S3, Table 4). In contrast, the logit proportion of *Lactobacillus* (5.8%), also a *Firmicutes* genus, did not vary with any of the fixed factors. The logit proportion of the second most abundant genus, *Actinomyces* (10.8%, *Actinobacteria*), differed unlike its phylum between seasons (*P* = 0.05), with higher relative abundances in summer, but did not differ by sex nor by other factors. *Varibaculum* (7.7%, also *Actinobacteria*) did not vary with any factor. The logit proportion of the third *Actinobacteria* genus, an uncultured *Coriobacteriales* genus (5.3%), was higher in males (*P* = 0.02), and was positively correlated with haptoglobin concentrations that were corrected for redness (*P* = 0.03, aviary contributed significantly to the model). The logit proportion of *Bacteroides* (5.6%, *Bacteroidetes*) varied similarly to its phylum with season*sex (*P* = 0.01), being higher in summer and females, and was positively correlated with BMR (*P* = 0.03). Lastly, the logit proportions of both *Fusobacteria* genera, *Oceanivirga* (5.8%) and *Fusobacterium* (5.3%) were similar to logit proportion of *Fusobacteria* higher in females (*P* = 0.01 and *P* = 0.04, respectively; note that aviary contributed significantly to the LMM for *Oceanivirga*). The logit proportion of *Oceanivirga* was also negatively correlated with residual haemolysis that had been corrected for sample age (*P* = 0.04).

### Bacterial biomarkers

LDA effect size analysis (LEfSe) detected 12 seasonal bacterial biomarkers in male homing pigeons, five of which were more abundant in winter and seven of which were more abundant in summer (Fig. 5a). The five winter LEfSe-based biomarkers in males belonged to the *Firmicutes* phylum, namely the most abundant ASV, *Lachnoclostridium 12* ASV nr. 093e1dd8072a68e5fa46226677183da, and its higher taxa up to class *Clostridia*. Of the seven summer LEfSe-based biomarkers in males, one belonged to *Firmicutes* (an *Enterococcus* ASV), two to *Actinobacteria* (*Actinomycetales* and *Actinomycetaceae*), two to *Bacteroidetes* (*Bacteroidetes* itself and *Bacteroidia*), and two to *Proteobacteria* (*Burkholderiaceae* and a *Ralstonia* ASV). In females, four seasonal biomarkers were present. One of which was more abundant in winter: a *Candidatus Arthromitus* ASV, belonging to the class *Clostridia* (*Firmicutes*). The three biomarkers that were more abundant in summer, belonged all to *Proteobacteria*: a *Pseudomonas* ASV, a *Delftia* ASV and the *Delftia* genus (Fig. 5b).

**Fig. 5.**
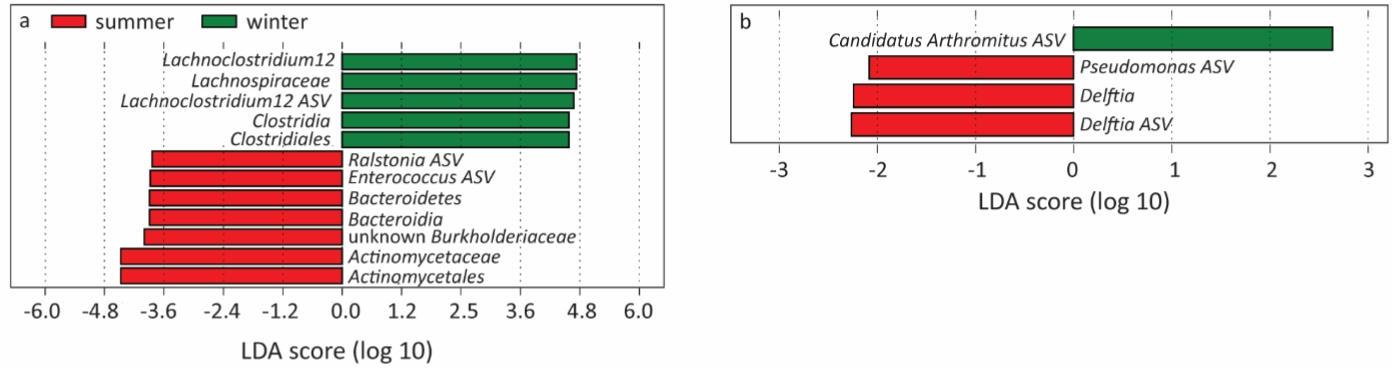
Seasonal LEfSe biomarkers per sex, and the significant LMM results in individual biomarkers. (**a**) Biomarkers that were more abundant in summer (negative values) or winter (–positive values) in male homing pigeons. (**b**) Biomarkers that were more abundant in summer or winter in female homing pigeons.

To determine biomarkers based on prevalence, we compared the summer and winter and female and male core microbiomes with the overall core microbiome, which was determined by taking all samples into account. The overall core microbiome (ASVs occurring in 90% of all samples) consisted of 13 ASVs, eight of which belonged to the *Actinobacteria* phylum and five to the *Firmicutes* phylum, indicating the importance of these phyla for homing pigeons (Table 5). Not surprisingly, the three most abundant ASVs belonged to the overall core microbiome: the *Lachnoclostridium 12* ASV (12.9%, *Firmicutes*), which was also a LEfSe-based winter biomarker in males, and a *Actinomyces* ASV (10.8%) and *Varibaculum* ASV (7.7%, both *Actinobacteria*). These ASVs occurred in all samples, as did a *Negativicoccus* ASV (4.1%, *Firmicutes*). The other core ASVs had intermediate abundances, but two core ASVs had low relative abundances: *Varibaculum* ASV (0.6%, *Actinobacteria*) and the unclassified *Propionibacteriaceae* ASV (0.3%, *Actinobacteria*).

**Table 5.**
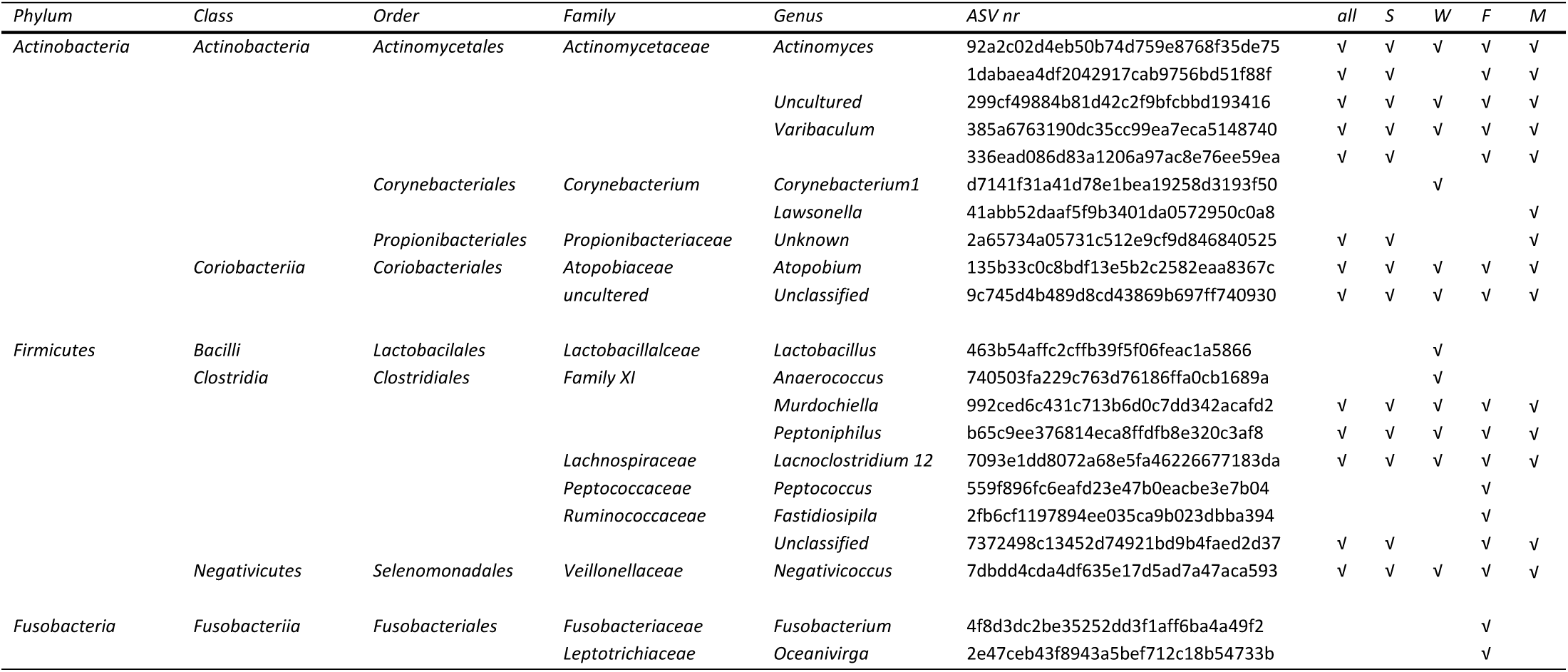
Core microbiome for all (all), summer (S), winter (W), female (F) or male (M) samples.

The summer core microbiome consisted of the same 13 ASVs as the overall core microbiome (Table 5), and thus we could not detect summer-specific bacterial biomarkers based on prevalence. However, three of the 12 winter core microbiome ASVs were unique to the winter core: a *Corynebacterium 1* ASV (*Actinobacteria*), and a *Lactobacillus* and *Anaerococcus* ASV (both *Firmicutes*, Table 5). In addition, four of the overall core ASVs were not present in the winter core microbiome: an *Actinomyces*, a *Varibaculum*, and the unclassified *Propionibacteriaceae* ASV (*Actinobacteria*), and an unclassified *Ruminococcaceae* ASV (*Firmicutes*).

Lastly, we determined core-based biomarkers for each sex. The male core microbiome consisted of 14 ASVs (Table 5), including the overall core microbiome plus one unique ASV, a *Lawsonella* ASV (*Actinobacteria*). The female core microbiome consisted of 16 ASVs, including four unique ASVs of which two belonged to the *Fusobacteria*, a phylum not present in the overall core microbiome, namely a *Oceanivirga* and a *Fusobacterium* ASV. The other two unique female core ASVs belonged to the *Firmicutes*, a *Fastidiosipila* and a *Peptococcus* ASV (Table 5). One ASV present in the overall core microbiome did not occur in the female core microbiome: the unclassified *Propionibacteriaceae* ASV (*Actinobacteria*) that was also not present in the winter core microbiome.

### Functional profile

The PICRUSt2 analysis yielded 161 KO metagenome functions in total. KO function abundances differed by season (PERMANOVA, *R*^2^ = 0.035, *F*_1, 53_ = 2.00, *P* = 0.03) and sex (*R*^2^ = 0.075, *F*_1, 53_ = 4.30, *P* = 0.03; Fig. 6a). The season*sex interaction was not significant. In females, three KO functions were more abundant in winter, and one KO function was more abundant in summer (LEfSe analyses; Fig. 6b). In males, 10 KO functions were more abundant in winter, and 11 KO functions were more abundant in summer (Fig. 6c). Males and females shared two KO functions that were more abundant in winter, while none of the summer-specific KO functions were shared. The shared winter-specific KO functions are important to metabolism and lipid metabolism: fatty acid biosynthesis (KO00061) and linoleic acid metabolism (KO00591). In addition, in males another winter-specific KO function is important to metabolism and lipid metabolism: KO01040 (biosynthesis of unsaturated fatty acids). In females, the third winter-specific KO function (KO00120, primary bile acid biosynthesis), may be involved in the response of gut microbiota to day length differences [28].

**Fig. 6.**
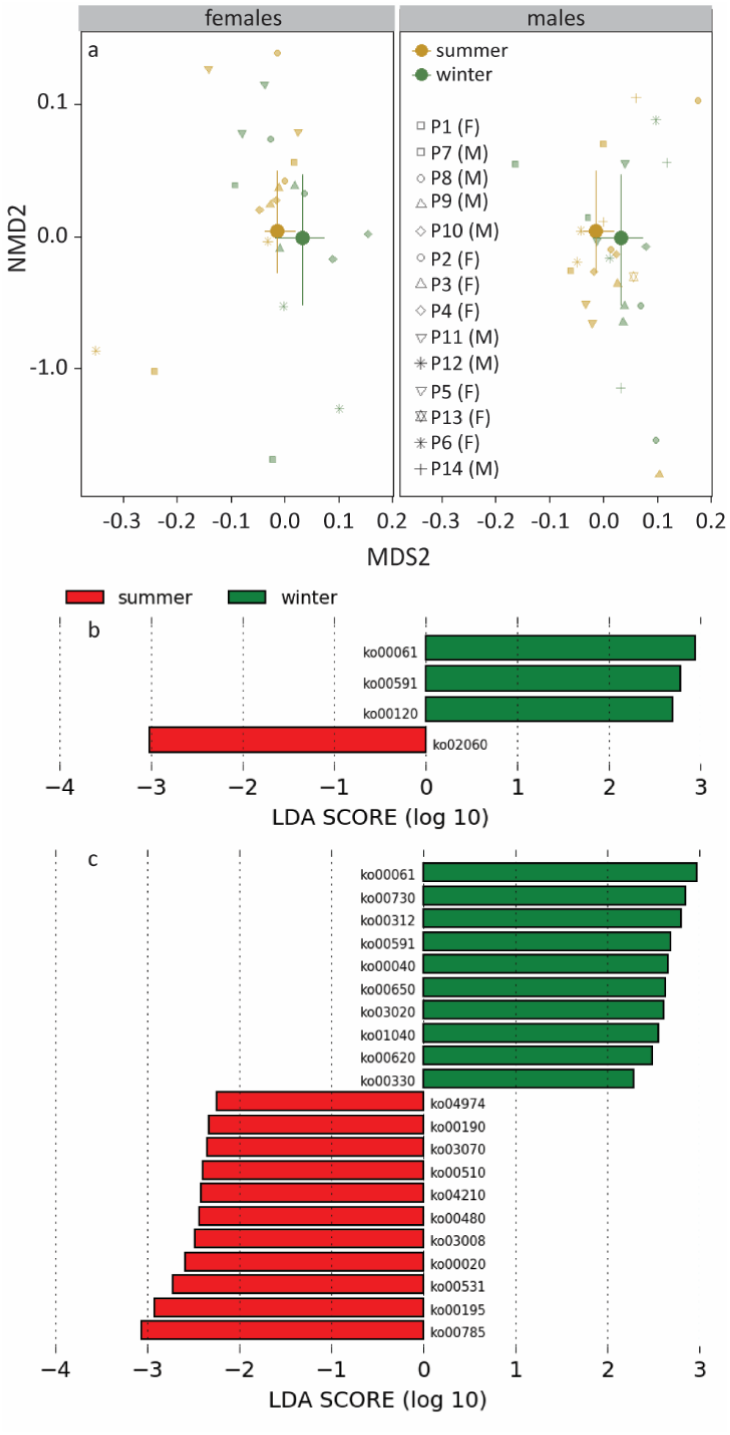
Seasonal and sexual variation in KO pathway abundances. (**a**) MDS plot of seasonal differences within sex. (**b**) KO pathways that were more abundant in summer or winter in females. (**c**) KO pathways that were abundant in summer or winter in males. Statistics are presented in the text. The large symbols represent the medians, the error bars the 25% and 75% quantiles. The transparent symbols present the underlying data for the individual pigeons. F, females, M, males.

## Discussion

We examined if other factors than diet, such as temperature and day length, play a role in shaping seasonal variation in gut microbiota in birds. Specifically, we investigated whether the gut microbiota of homing pigeons that lived outdoors differed between summer and winter despite a constant diet. We tested whether seasonal variation in gut microbiota was correlated with host metabolism, immune function of both. Metabolism is a potential intermediary between seasonal changes in temperature and the animal’s gut microbiota. Similarly, immune competence might link seasonal changes in day length and the gut microbiota. All gut microbiota characteristics showed summer-winter differences (expectation 1). Temperature likely contributed to the summer-winter differences in gut microbiota, as the relative abundances of *Firmicutes* tended to be higher in winter and relative abundances of *Bacteroidetes* were higher in summer (expectation 2), and multiple gut microbiota characteristics were correlated with at least one metabolism index (expectation 3). Lastly, we found correlations between immune indices and gut microbiota characteristics (expectation 4).

In addition to summer-winter differences, most gut microbiota indices differed between males and females. These sex difference may have been driven by corresponding differences in diet. For example, we documented that the percentage pellets in the diet was 1.8 times higher in females than in males. However, the pellets and seed mixture did not differ much in terms of nutrition (i.e., crude protein, crude fat, crude fibre, crude ash contents, and energy content; Table S1). Instead, sex-specific physiological mechanisms may have played a more important role in structuring gut microbiota within each sex, but the nature and consequences of such mechanisms require further investigation.

### Seasonal temperature variation contributed to seasonal gut microbiota variation

A strong indication that seasonal temperature differences partly caused the summer-winter differences in the gut microbiota was the tendency of the relative abundances of *Firmicutes*, a phylum previously associated with low temperature [20, 23], to peak in winter, and the many *Firmicutes* taxa present among the winter bacterial biomarkers. The higher winter relative abundances of *Firmicutes* mainly resulted from changes in the *Clostridia* class, which contained the most abundant genus (*Lachnoclostridium 12*) and ASV (*Lachnoclostridium 12* ASV). These *Lachnoclostridium 12* taxa were LEfSe-based winter biomarkers in males, and peaked over all samples in abundance in winter just as their higher taxa (*Lachnospiraceae, Clostridiales, Clostridia*). These taxa are thus important components of the winter gut microbiota in homing pigeons. In humans, *Lachnospiraceae* are recognized as an essential part of the core microbiome that promotes health [62]. *Lachnospiraceae* comprises anaerobic, fermentative, and chemoorganotrophic bacteria, that produce short-chain fatty acids (SCFAs) like butyrate by hydrolysing carbohydrates [62]. SCFAs fulfill vital functions in animals. They provide an energy source, maintain intestinal epithelium physiology, regulate innate and adaptive immune function, and may reduce inflammation [63–66], but SFCAs may also influence the regulation and capacity of energy regulation [67]. Of the two KO functions that were more abundant in winter in both sexes, one (KO00061) represents fatty acid biosynthesis and the other (KO00591) linoleic acid metabolism. Considering the importance of SCFAs, their increased biosynthesis by the gut microbiota may be especially beneficial. SCFAs produced by gut microbiota might even contribute to overwinter survival of hosts. In winter, energy budges can come under pressure, for example, due to increased thermoregulation and foraging costs, but gut microbiota-produces SCFAs may alleviate some of this pressure. Enhanced bacterial metabolism of linoleic acid, a polyunsaturated omega-6 fatty acid (PUFA; 18:2*n*6), may also offer advantages. High-PUFA diets are beneficial to migrating birds because they reduce the energy expenditure during long-duration flights, which are otherwise energetically demanding [67,68]. This benefit may be due to PUFAs increasing the amount of transport proteins and catabolic enzymes that deliver fatty acids to mitochondria [69]. Linoleic acid and other PUFAs may offer similar benefits to wintering birds facing increased energy expenditures.

Our results showed agreements and disagreements with 18 published studies on the effects of temperature on gut microbiota in vertebrates (mammals, birds, reptiles, amphibians, and fish; Fig. 7 and Supplementary information Table S3). Most of these studies focused directly on temperature effects (14 lab studies), but some focussed on seasonal effects (one husbandry and three field studies). In a majority of the studies, including ours, relative abundances of *Firmicutes* were highest at lower temperatures and relative abundances of *Bacteroidetes* were higher at higher temperatures. *Bacteroidetes* also ferment carbohydrates and produce SCFAs [70]. The alternating peaks in relative abundances of *Firmicutes* and *Bacteroidetes* between seasons, suggests that differences in the carbohydrate fermentation products between these taxa may play a role in seasonal acclimatization in homing pigeons and other vertebrates. The alternating peak also resulted in a tendency of the *Firmicutes*:*Bacteroidetes* ratio to be higher in winter, a result that was most evident in males (Fig S4). In mammals, the cold-associated increase in the *Firmicutes*:*Bacteroidetes* ratio is associated with aspects of cold acclimatization in host metabolism. The higher ratio is associated with enhanced energy extraction and thus increased energy consumption [23]; it is also associated with high-fat diets [71] and obesity [72]. Additional body mass in winter, as we observed in our homing pigeons, is an adaptive trait in animals living in temperate or cold areas. Increased body reserves promote survival during the harsher winter period. A decrease in the *Firmicutes*:*Bacteroidetes* ratio at warmer temperatures is associated with fasting [71] and protection against obesity [72]. These affects, especially the latter, are beneficial in summer in wild animals. In addition, it is noteworthy that both *Firmicutes* and *Bacteroidetes* relative abundances were correlated with body mass or BMR, confirming that also in homing pigeons, host metabolism may mediate the temperature effects on gut microbiota. All in all, in homing pigeons the observed seasonal variation in the *Firmicutes*:*Bacteroidetes* ratio suggests that seasonal patterns in gut microbiota can be attributed to acclimatization to seasonal temperature changes.

**Fig. 7.**
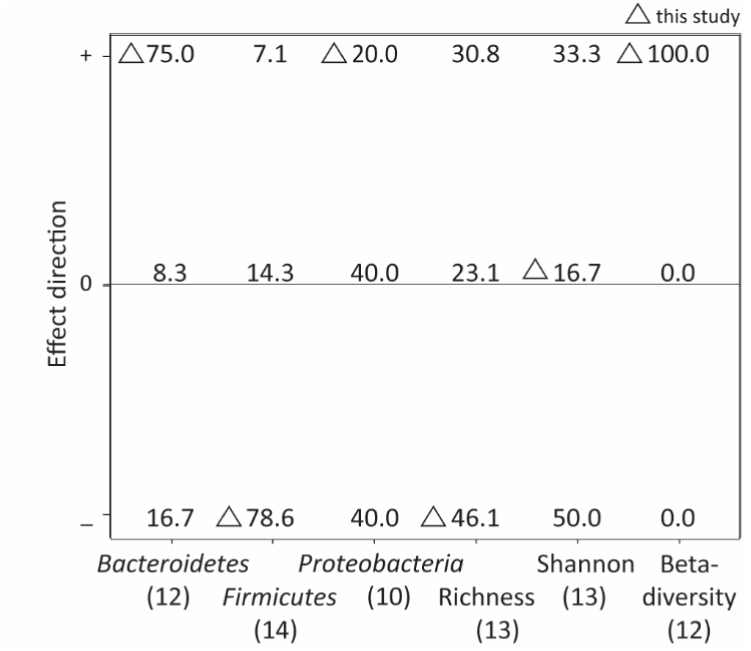
The effect directions of temperature on some aspects of vertebrate gut microbiota found in 18 studies and 17 species. Literature sources and data, including some additional data, are presented in Supplementary information Table S3. A positive effect direction (+) indicates an increase with temperature or a significant difference in beta-diversity, a negative effect direction (–) indicates a decrease with temperature (does not apply to beta-diversity), and the 0 indicates no significant effect. The numbers indicate the percentage of studies with that effect direction within the studies that reported on the concerned index. The number of studies that presented data for the given taxa, alpha- and beta-diversity indices are indicated between brackets after the X-axe labels.

Many of the 18 studies also reported the effects of temperature on alpha-diversity indices (richness and Shannon index) and the relative abundances of *Proteobacteria* (Fig. 7). There was no general trend for these variables. Beta-diversity indices, on the other hand, always differed between temperatures or seasons when reported, similar to this study. Thus, our results closely match the general temperature effects on gut microbiota presented in the literature, indicating that the seasonal variation in environmental temperature contributed considerably to the summer-winter differences in the gut microbiota of homing pigeons.

### Is immune competence a link between day length and gut microbiota?

The gut microbiota is known to have strong and dynamic interactions with especially the innate immune system [64–66]. For instance, the SFCAs produced by the gut bacteria play an essential role in the host (intestinal) immune defence. These molecules interact with the intestinal epithelial cells, reduce intestinal inflammation, provide protection against pathogens, and regulate activation and differentiation of immune cells [64–66]. We found multiple correlations between the immune indices and gut microbiota characteristics, but not between immune indices and the beta diversity indices. However, these correlations did not reveal consistent involvement of one or more specific immune indices. This lack of consistency complicates interpretations. Moreover, in contrast to the metabolism indices, only two of the seven immune function indices (i.e., antibody titres to PC-BSA and haptoglobin concentration corrected for redness) showed seasonal variation. Of these, only the haptoglobin concentration was correlated with an aspect of the gut microbiota (relative abundances of uncultured *Coriobacteriales*). Given this complexity of the results and lack of seasonal differences in the immune indices, our study does not clearly support the idea that innate immune function indices mediates the previously documented links between daylength and gut microbiota. The lack of clear correlations between the gut microbiota and the innate immune indices may also be due to the modest number of individuals included in the study (14 homing pigeons), because generally, and also here, there is a large individual variation in host-associated microbiota. Note that we nevertheless do find consistent and strong effects of temperature to the seasonal variation in gut microbiota.

## Conclusions

Seasonal environmental variation influenced the gut microbiota in homing pigeons, even when the birds were fed a constant diet. Temperature likely drove part of the seasonal differences in the gut microbiota composition because we found multiple correlations between gut microbiota characteristics and metabolism indices. Furthermore, the summer-winter differences in gut microbiota characteristics matched previously described effects of temperature variation on vertebrate gut microbiota. In addition, in winter, the *Firmicutes*:*Bacteroidetes* ratio tended to be higher and fatty acid related KO functions were more abundant, indicating that the seasonal variation in gut microbiota contributes to seasonal acclimatization of the host. We found less consistent correlations between gut microbiota characteristics and innate immune indices, and we conclude conservatively that the here used innate immune competence may be an unlikely link between day length and the gut microbiota. However, we do not exclude that day length may have contributed to the seasonal differences in gut microbiota. Overall, our results highlight the need for future studies that disentangle different seasonally-varying factors (i.e., temperature, daylength, behaviour, diet, etc.) if the goal is to fully understand the causal mechanisms driving seasonal variation in gut microbiota.

## Acknowledgements

We thank Henri Zomer for doing the DNA extractions and Yiran Sun, and Thirsa van Wichen for helping perform the ELISA analyses. We thank Pieter van Veelen and Stephanie Vink for their advice on the data analysis and Pieter for his advice on the lab work. Our Centre for Information Technology provided support and access to the Peregrine high-performance computing cluster. This work was supported by the NWO-Vidi grant 864.10.012 to BIT.

## Authors’ contributions

MWD, KDM, MAV, and BIT designed and performed the experiment. KDM, JAJA, HKP, MAV, and MvdV performed the lab analyses. MvdV processed the sequence data with QIIME2 and PICRUSt. MWD did the statistical analysis in R. MAV, JFS and BIT advised on the statistical analyses. MWD wrote the original draft. All authors contributed to writing, reviewing and editing the text.

## Ethics approval

This study was performed under animal welfare license no. 5095 of the Institutional Animal Care and Use Committee of the University of Groningen, complying with the Dutch and European Laws.

## Competing interests

The authors declare that they have no competing interests.

## Notes

### Competing Interest Statement

The authors have declared no competing interest.

