## Supplementary information for "Gut microbiota of homing pigeons shows summer-winter variation under constant diet indicating a substantial effect of temperature"

### Contents **Page**

**Calculation of the antibody titres** 2
**Table S1** Compositions of the two food types offered to the homing pigeons 3
**Table S2** Overview of the KEGG pathways (KOs) specific for winter and summer per sex 3
**Table S3** Ambient temperature effects on vertebrate gut microbiota: phylum relative abundance, alpha-diversity, and beta-diversity 4
**Fig. S1** Richness rarefaction curves 5
**Fig. S2** Results of the Procrustes analyses 5
**Fig. S3** Significant variation in the Logit-transformed proportions of the most abundant genera with season, sex, metabolism or immune indices 8
**Fig S4** Seasonal variation in the ratio *Firmicutes*/*Bacteroidetes* proportions in (**a**) females and (**b**) males. 9

### Calculation of the antibody titres

Antibody titres were calculated as described by [1] (taken from [2]). Briefly, the optical densities (OD) of the duplicate standard positive plasma were averaged for each plate. Logit values of the OD per plate were calculated using:

$$Logit\ OD = ln\ (OD/(OD_{max} - OD))$$

Where OD is the OD of a well, and $OD_{max}$ is the maximum averaged OD of the duplicate standard positive plasma samples. The last positive well (lpw) of the averaged duplicate standard positive plasma sample was set to the sixth dilution. A linear regression line of the logit OD against the respective $log_2$-dilution values of the averaged duplicate standard positive plasma samples was determined, which resulted in a regression coefficient β. Titres of the plasma samples per plate were calculated using:

$$Titre = (logit\ OD_{lpw} - (logit\ OD_{sample} - \beta\ x\ log_2(dilution_{sample})))/\ \beta$$

Were logit $OD_{lpw}$ is the estimated logit OD at the lpw calculated with the estimated linear regression function using the $log_2$-dilution value of that well, logit $OD_{sample}$ is the logit OD calculated of the OD closest to 50% of $OD_{max}$ for a plasma sample of an individual ($OD_{sample}$), β is the regression coefficient of the estimated linear regression function of the averaged duplicate standard positive plasma samples, and $log_2$ ($dilution_{sample}$) is the $log_2$-dilution value at which $OD_{sample}$ occurred.

**Table S1** Compositions of the two food types offered to the homing pigeons.

| | *4 seasons Kasper™ 6705* | *Pellets Kasper™ P40* |
|---|---|---|
| Crude protein | 12.3% | 15.5% |
| Crude fat | 3.0% | 2.8% |
| Crude fibre | 3.4% | 2.2% |
| Crude ash | 1.8% | 4.0% |
| Composition | Corn, wheat, sorghum, yellow peas, barley, dunpeas, green peas, white sorghum, safflower seed, peeled oats and soybean oil. | Corn, wheat, soybean meal feed, corn gluten, calcium carbonate, monocalcium phosphate, dextrose, sodium chloride, alfalfa, timothee grassmeal, grassmeal, sunflower seed extracted, dried beet pulp, cane molasses, rape seed extracted, linseed, linseed expeller, soybean oil, and added vitamins, minerals, and amino acids. |

**Table S2** Overview of KEGG ortholog (KO) functions specific for winter and summer per sex.

| Sex | KO winter | Description | KO summer | Description |
|---|---|---|---|---|
| **Males** | KO00040 | Pentose and glucuronate interconversions | KO00020 | Citrate cycle (TCA cycle) |
| | | | KO00190 | Oxidative phosphorylation |
| | KO00061 | Fatty acid biosynthesis | KO00195 | Photosynthesis |
| | KO00312 | Beta-Lactam resistance | KO00480 | Glutathione metabolism |
| | KO00330 | Arginine and proline metabolism | KO00510 | N-Glycan biosynthesis |
| | KO00591 | Linoleic acid metabolism | KO00531 | Glycosaminoglycan degradation |
| | KO00620 | Pyruvate metabolism | KO00785 | Lipoic acid metabolism |
| | KO00650 | Butanoate metabolism | KO03008 | Ribosome biogenesis in eukaryotes |
| | KO00730 | Thiamine metabolism | KO03070 | Bacterial secretion system |
| | KO01040 | Biosynthesis of unsaturated fatty acids | KO04210 | Apoptosis |
| | | | KO04974 | Protein digestion and absorption |
| | KO03020 | RNA polymerase | | |
| **Females** | KO00061 | Fatty acid biosynthesis | KO02060 | Phosphotransferase system (PTS) |
| | KO00120 | Primary bile acid biosynthesis | | |
| | KO00591 | Linoleic acid metabolism | | |

**Table S3** Ambient temperature effects on vertebrate gut microbiota: phylum relative abundance, alpha-diversity, and beta-diversity.

| *Species* | *Location* | *Cold exposure* | *Microbiota effects* [a] | *Source* |
|---|---|---|---|---|
| <u>Mammals</u> | | | | |
| Alpine musk deer (*Moschus chrysogaster*) | Breeding centre,at Xinglong Mountain. China | Natural temperatures in spring and winter | *Firmicutes* ↘<br>*Bacteroidetes* ↗<br>Richness ↘<br>Shannon ↘<br>Beta-diversity + | [1] |
| Brandt's vole (*Lasiopodomys bandtii*) | Lab | 3 weeks at 4°C or 23°C | *Firmicutes* –<br>*Bacteroidetes* ↗<br>*Proteobacteria* –<br>Faith's PD –<br>Beta-diversity + | [2] |
| Brandt's vole (*Lasiopodomys bandtii*) | Lab | 4 weeks at 4°C, 4 weeks at 4°C followed by 4 weeks at 23°C, or 4-8 weeks at 23°C | *Firmicutes* ↘<br>*Bacteroidetes* ↗<br>*Proteobacteria* –<br>Richness ↘<br>Shannon ↘ | [3] |
| Forest musk deer (*Mochus berezovskii*) | Breeding centre, at the Qinghai-Tibet plateau, China | Natural temperatures in summer and winter | *Firmicutes* ↘<br>*Bacteroidetes* ↗<br>Richness ↘<br>Shannon ↘<br>Beta-diversity + | [1] |
| Mouse C57Bl/6J (*Mus musculus*) | Lab | Up to 10 days at 6°C or room temperature | *Firmicutes* ↘<br>*Bacteroidetes* ↗<br>*Proteobacteria* –<br>*Verrucomicrobia* ↗ | [4] |
| Mouse C57BL6/J (*Mus musculus*) | Lab | Up to 6 days at 12°, 17° or 23°C | *Firmicutes* ↘<br>*Bacteroidetes* ↗<br>*Proteobacteria* ↗<br>Faith's PD ↗ | [5] |
| Mouse wild-type (*Mus musculus*) | Lab | 6 days at 6°C or 30°C | Richness ↗<br>Shannon ↗<br>Beta-diversity + | [6] |
| Siberian flying squirrel (*Pteromys volans orii*) | Field: Hokkaido forest, Japan | May-August, 8°-22°C | Beta-diversity + | [7] |
| Wild blue sheep (*Pseudois nayaur*) | Field: Helan Mountain, China | Summer (~17°C) or winter (~-9°C) | *Firmicutes* ↗<br>*Bacteroidetes* –<br>Shannon + | [8] |
| <u>Birds</u> | | | | |
| Greater sage-grouse (*Centrocercus urophasianus*) | Field: Sublette (S) & Natrona (N) county, WY, USA | September (S) and December (S & N), at 20°C or -3°C | Richness ↗<br>Shannon ↗<br>Beta-diversity + | [9] |
| Layer (*Gallus gallus domesticus*) | Commercial husbandry, Dafreng, Jiansu, China | May (min-max: 20.8°-25.4°C) or July (min-max: 28.6°-31.8°C) | *Firmicutes* ↘<br>*Bacteroidetes* ↗<br>*Proteobacteria* ↘<br>*Fusobacteria* ↘ | [10] |
| <u>Reptiles</u> | | | | |
| Common lizard (*Zootoca vivipara*) | Lab | Present (26.6°C), intermediate (28.2°C) or warm (28.4°C) for 3 | *Firmicutes* ↘<br>*Bacteroidetes* ↘<br>*Proteobacteria* ↗ | [11] |

| | | | | |
|---|---|---|---|---|
| | | months, sampled 8 months later | *Actinobacteria* ↗<br>Richness ↘<br>Beta-diversity + | |
| Western fence lizard (*Sceloporus occidentalis*) | Wild-caught in lab | Control at 25°C, experimental moved to 35°C after 7d at 25°C | *Firmicutes* ↘<br>Richness −<br>Shannon −<br>Beta-diversity + | [12] |
| <u>Amphibians</u> | | | | |
| Eastern red-backed salamander (*Plethodon cinereus*) | Lab | Six days at 10°C, 15°C and 20°C | Richness ↘<br>Shannon ↘<br>Faith's PD ↘ | [13] |
| Northern leopard frog (*Lithobates pipiens*) tadpoles | Lab | Raised at 18°C or 28°C, samples at Gosner stage 38.3. | *Firmicutes* ↘<br>*Bacteroidetes* −<br>*Proteobacteria* ↘<br>*Planctomycetes* ↗<br>*TM6* ↗<br>Alpha-diversity −<br>Beta-diversity + | [14] |
| Western clawed frog (*Xenopus tropicalis*) | Lab | Raised at low (23°C) or high (28°C) temperatures, sampled as froglets, 3 days after metamorphosis | *Firmicutes* ↘<br>*Bacteroidetes* ↗<br>*Proteobacteria* ↘<br>*Verrucomicrobia* ↗<br>Richness −<br>Shannon −<br>Weighted UNIF +<br>Unweighted UNIF − | [15] |
| <u>Fish</u> | | | | |
| Blue tilapia (*Oreochromis aureus*) | Lab | Lines selected for cold tolerance (12°C) or not (control). Two days at 12°C or 24°C | *Firmicutes* −<br>*Bacteroidetes* ↗<br>*Proteobacteria* −<br>*Planctomycetes* ↗<br>*Verrucomicrobia* ↗<br>*TM6* ↘<br>Richness ↗<br>Shannon ↗<br>Beta-diversity + | [16] |
| Milkfish (*Chanos chanos*) | Lab | Control (26°C) vs elevated temperature (33°C) at days 0, 14 and 21 [b]. | *Fusobacteria* ↗<br>*Proteobacteria* ↘<br>Simpson d0 ↘<br>Beta-diversity + | [17] |
| Rainbow trout (*Oncorhynchus mykiss*) | Lab | 11°C vs 18°C, sampled after a week | *Firmicutes* ↘<br>Richness ↘<br>Shannon ↘ | [18] |
| Yellowtail kingfish juveniles (*Seriola lalandi*) | Lab | Control 24°C vs 20°C and 26°C after 30 days | Richness ↗<br>Shannon ↘ | [19] |

[a] The microbiota effects included were selected as follows: 1) the phyla *Firmicutes* and *Bacteroidetes* were included as they are generally reported to vary with temperature or season, 2) other phyla included in the analyses, were included in the table 3) sometimes effect directions in alpha-diversity varied between measures used, hence we report the measure type used, and 4) this differentiation did not occur in the beta-diversity, hence we did not differentiate here. Symbols: ↘, decrease with temperature; ↗, increase with temperature; −, no effect; +, significant difference. Abbreviation: PD, phylogenetic diversity. [b] The largest effect of a temperature increase was found at day 0, i.e., after the week required to increase the temperature from 26°C to 33°C (increased by 1°C per day).

**References to Table S3**

1. Jiang F, Gao H, Qin W, Song P, Wang H, Zhang J, et al. Marked seasonal variation in structure and function of gut microbiota in forest and alpine musk deer. Front Microbiol. 2021;12:1–13.
2. Zhang X-Y, Sukhchuluun G, Bo T-B, Chi Q-S, Yang J-J, Chen B, et al. Huddling remodels gut microbiota to reduce energy requirements in a small mammal species during cold exposure. Microbiome. 2018;6:103.
3. Bo TB, Zhang XY, Wen J, Deng K, Qin XW, Wang DH. The microbiota–gut–brain interaction in regulating host metabolic adaptation to cold in male Brandt's voles (*Lasiopodomys brandtii)*. ISME J. 2019;13:3037–53.
4. Chevalier C, Stojanović O, Colin DJ, Suarez-Zamorano N, Tarallo V, Veyrat-Durebex C, et al. Gut microbiota orchestrates energy homeostasis during cold. Cell. 2015;163:1360–74.
5. Ziętak M, Kovatcheva-Datchary P, Markiewicz LH, Ståhlman M, Kozak LP, Bäckhed F. Altered microbiota contributes to reduced diet-induced obesity upon cold exposure. Cell Metab. 2016;23:1216–23.
6. Worthmann A, John C, Rühlemann MC, Baguhl M, Heinsen FA, Schaltenberg N, et al. Cold-induced conversion of cholesterol to bile acids in mice shapes the gut microbiome and promotes adaptive thermogenesis. Nat Med. 2017;23:839–49.
7. Liu PY, Cheng AC, Huang SW, Chang HW, Oshida T, Yu HT. Variations in gut microbiota of Siberian flying squirrels correspond to seasonal phenological changes in their Hokkaido subarctic forest ecosystem. Microb Ecol. 2019;78:223–31.
8. Zhu Z, Sun Y, Zhu F, Liu Z, Pan R, Teng L. Seasonal variation and sexual dimorphism of the microbiota in wild blue sheep (*Pseudois nayaur*). Front Microbiol. 2020;11:1260.
9. Graves GR, Schmidt BK, O'Mahoney MJ V., Matterson KO, Drovetski S V. Distinct microbiotas of anatomical gut regions display idiosyncratic seasonal variation in an avian folivore. Anim Microbiome. 2019;1:1–11.
10. Zhu L, Liao R, Wu N, Zhu G, Yang C. Heat stress mediates changes in fecal microbiome and functional pathways of laying hens. Appl Microbiol Biotechnol. 2019;103:461–72.
11. Bestion E, Jacob S, Zinger L, Di Gesu L, Richard M, White J, et al. Climate warming reduces gut microbiota diversity in a vertebrate ectotherm. Nat Ecol Evol. 2017;1:1–3.
12. Moeller AH, Ivey K, Cornwall MB, Herr K, Rede J, Taylor EN, et al. Lizard gut microbiome changes with temperature and is associated with heat tolerance. Appl Environ Microbiol. 2020;
13. Fontaine SS, Novarro AJ, Kohl KD. Environmental temperature alters the digestive performance and gut microbiota of a terrestrial amphibian. J Exp Biol. 2018;221:jeb.187559.
14. Kohl KD, Yahn J. Effects of environmental temperature on the gut microbial communities of tadpoles. Environ Microbiol. 2016;18:1561–5.
15. Li J, Rui J, Li Y, Tang N, Zhan S, Jiang J, et al. Ambient temperature alters body size and gut microbiota of *Xenopus tropicalis*. Sci China Life Sci. 2020;63:915–25.
16. Kokou F, Sasson G, Nitzan T, Doron-Faigenboim A, Harpaz S, Cnaani A, et al. Host genetic selection for cold tolerance shapes microbiome composition and modulates its response to temperature. Elife. 2018;7:1–21.
17. Hassenrück C, Reinwald H, Kunzmann A, Tiedemann I, Gärdes A. Effects of thermal stress on the gut microbiome of juvenile milkfish (*Chanos chanos*). Microorganisms. 2021;9:1–18.
18. Huyben D, Sun L, Moccia R, Kiessling A, Dicksved J, Lundh T. Dietary live yeast and increased water temperature influence the gut microbiota of rainbow trout. J Appl Microbiol. 2018;124:1377–92.
19. Soriano EL, Ramírez DT, Araujo DR, Gómez-Gil B, Castro LI, Sánchez CG. Effect of temperature and dietary lipid proportion on gut microbiota in yellowtail kingfish Seriola lalandi juveniles. Aquaculture. 2018;497:269–77.

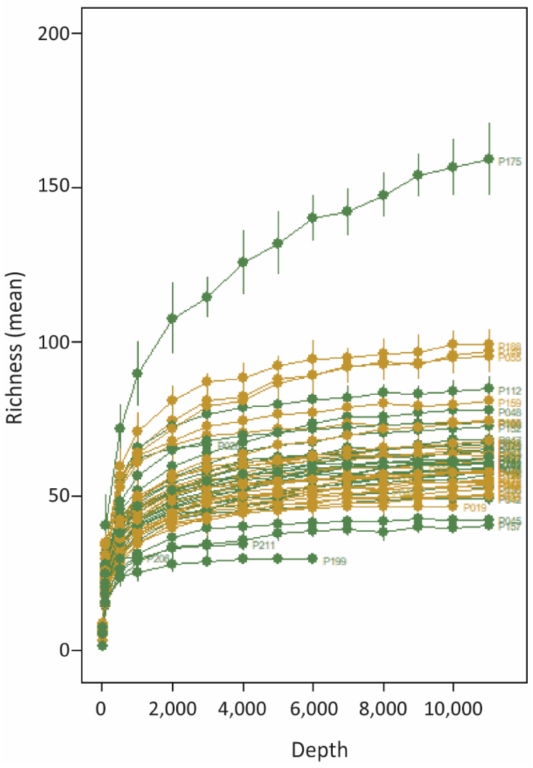


**Fig. S1** Richness rarefaction curves. Most curves levelled off around 3,000 reads.

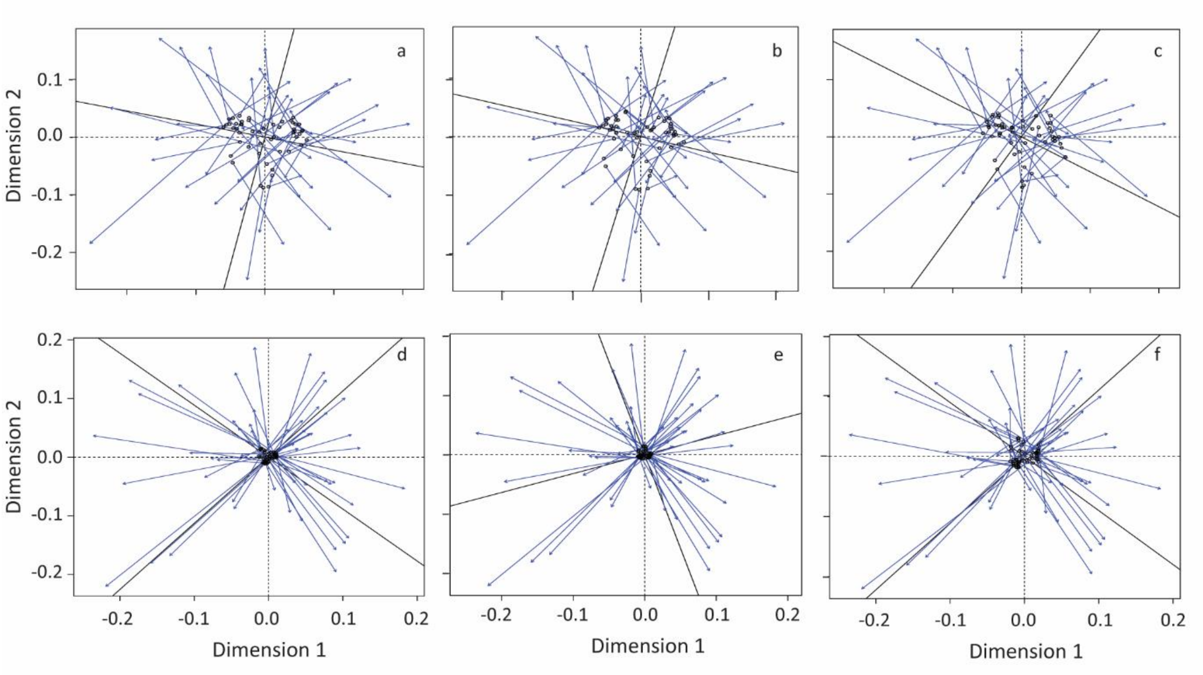


**Fig. S2** Results of the Procrustes analyses. The ordination dimensions of the metabolism indices versus the ordination dimensions of the Jaccard (**a**), Brays-Curtis (**b**) and weighted UniFrac (dis)similarities and distances (**c**). And the ordination dimensions of the seven innate immune indices versus the ordination dimensions of the Jaccard (**d**), Brays-Curtis (**e**) and weighted UniFrac (**f**).

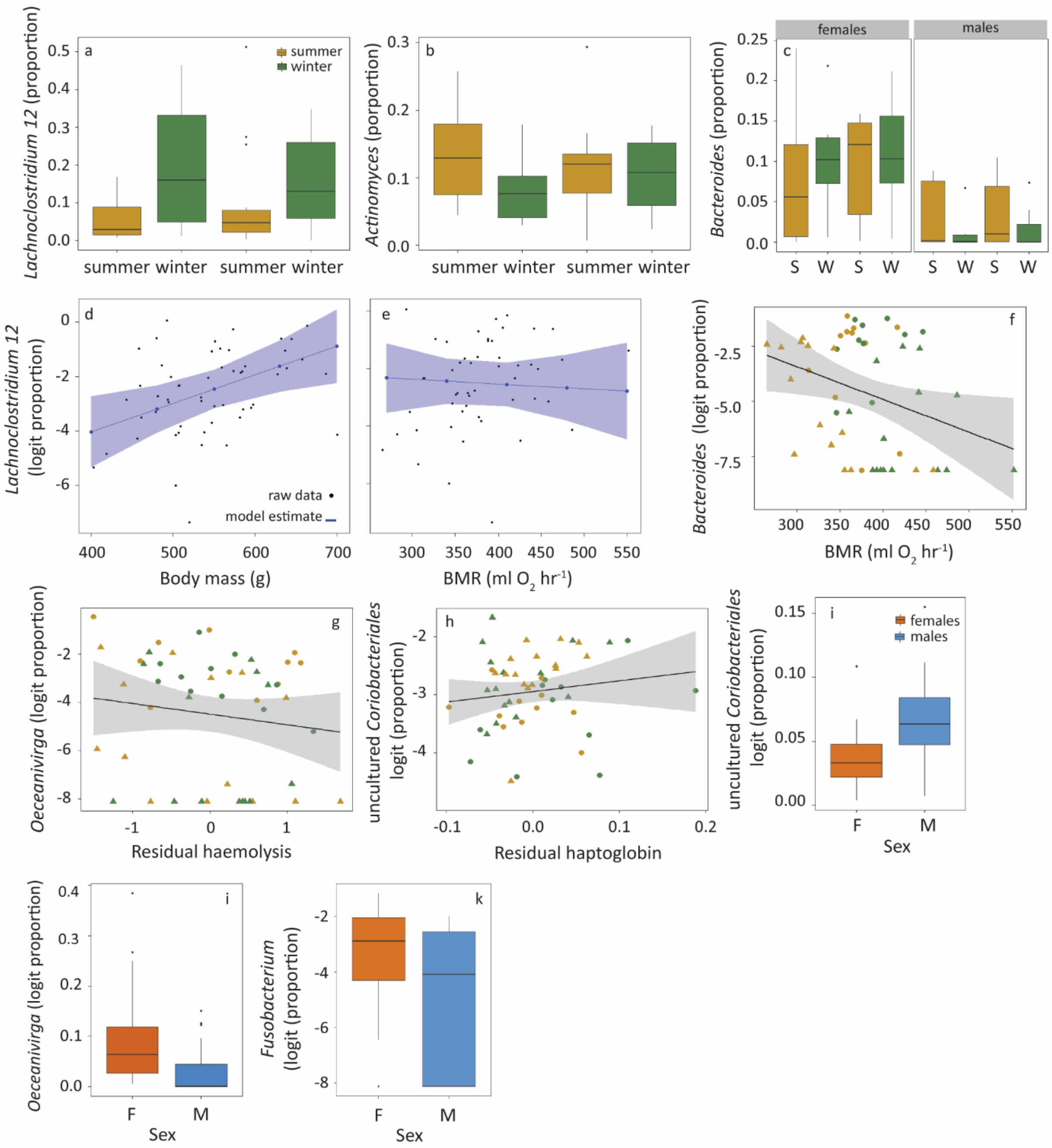


**Fig. S3** Significant variation in the Logit-transformed proportions of the most abundant genera with season, sex, metabolism or immune indices. Boxplots of the seasonal variation in *Lachnoclostridum12* (**a**), *Actinomyces* (**b**), and *Bacteroides* (**c**), for the latter per sex. Seasons (S = summer, W = winter) start with the summer of 2013. The model estimates for the linear relationships between *Lachnoclostridum12* and BMR and body mass are presented in panels (**d**, **e**). Logit-transformed proportions of *Bacteroides* increased with increasing BMR (**f**). *Oceanivirga* decreased with residual haemolysis (**g**) and uncultured *Coriobacterium* increased with increasing residual haptoglobin (**h**). The uncultured *Coriobacterium*, *Oceanivirga* and *Fusobacterium* also varied with sex (**i**, **j**, **k**; F, females, M, males). Statistics are presented in Table 3 of the main text.

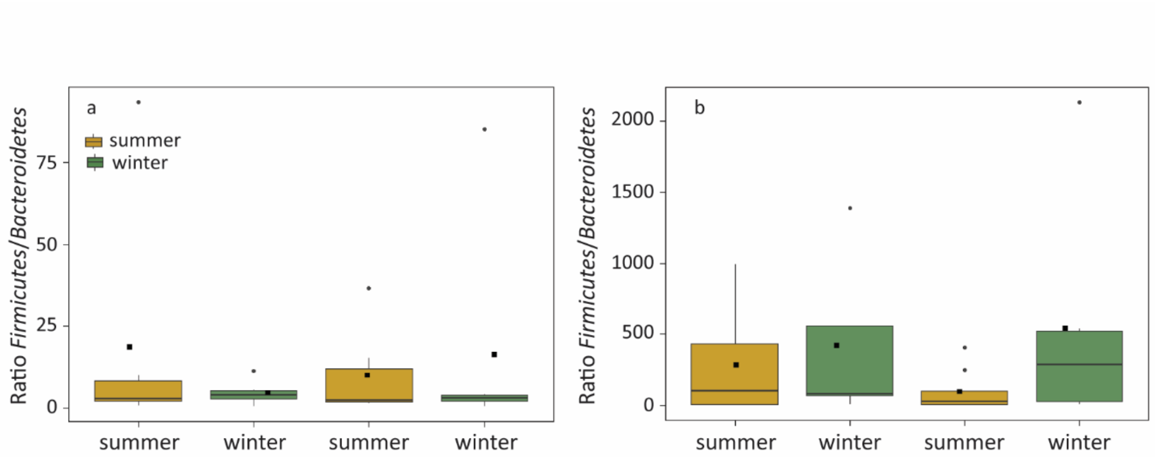


**Fig. S4** Seasonal variation in the *Firmicutes*:*Bacteroidetes* ratio in (**a**) females and (**b**) males. There was a significant effect of the interaction season*sex on the *Firmicutes*:*Bacteroidetes* ratio (LMM, $P$ = 0.04, aviary did contribute significantly to the model, $P$ = 0.01). Note the different ranges of the y-axes.
